# Performance and limitations of linkage-disequilibrium-based methods for inferring the genomic landscape of recombination and detecting hotspots: a simulation study

**DOI:** 10.1101/2022.03.30.486352

**Authors:** Marie Raynaud, Pierre-Alexandre Gagnaire, Nicolas Galtier

## Abstract

Knowledge of recombination rate variation along the genome provides important insights into genome and phenotypic evolution. Population genomic approaches offer an attractive way to infer the population-scaled recombination rate ρ=4*N*_*e*_*r* using the linkage disequilibrium information contained in DNA sequence polymorphism data. Such methods have been used in a broad range of plant and animal species to build genome-wide recombination maps. However, the reliability of these inferences has only been assessed under a restrictive set of conditions. Here, we evaluate the ability of one of the most widely used coalescent-based programs, *LDhelmet*, to infer a genomic landscape of recombination with the biological characteristics of a human-like landscape including hotspots. Using simulations, we specifically assessed the impact of methodological (sample size, phasing errors, block penalty) and evolutionary parameters (effective population size (*N*_*e*_), demographic history, mutation to recombination rate ratio) on inferred map quality. We report reasonably good correlations between simulated and inferred landscapes, but point to limitations when it comes to detecting recombination hotspots. False positive and false negative hotspots considerably confound fine-scale patterns of inferred recombination under a wide range of conditions, particularly when *N*_*e*_ is small and the mutation/recombination rate ratio is low, to the extent that maps inferred from populations sharing the same recombination landscape appear uncorrelated. We thus address a message of caution for the users of these approaches, at least for genomes with complex recombination landscapes such as in humans.

## Introduction

Recombination is highly conserved among sexually reproducing species of eukaryotes. This fundamental mechanism of meiosis is essential for the proper segregation of homologous chromosomes during the reductional division. Recombination involves crossing over events (CO) that play a crucial evolutionary role by allowing genetic mixing and generating new combinations of alleles (Baudat and de Massy 2007; Cromie et al. 2001; Capilla et al. 2016). Measuring the rate at which recombination occurs and the magnitude of its variation along the genome has important implications for fundamental research in molecular biology and evolution (Stapley et al. 2017), but also for applied genomics such as genome-wide association studies (GWAS) (Morris et al. 2013; Hunter et al. 2016). Several approaches have been developed to reconstruct genome-wide recombination maps (reviewed in Peñalba and Wolf 2020). Cytological methods, like ChIP-seq, target protein-DNA complexes directly involved in the formation of double-strand breaks (DSB) and CO during meiosis (Pratto et al. 2014). Gamete typing methods analyse the meiotic products of a diploid individual (reviewed in (Carrington and Cullen 2004; Dréau et al. 2019; Sun et al. 2019). Methods based on pedigree analysis reconstruct the gametic phase from patterns of allele inheritance in bi-parental crosses (Lander et Green 1987; Kong et al. 2002; Kodama et al. 2014; Rastas 2017). All these approaches have the advantage of providing direct estimates of the recombination rate. However, by focusing on CO that occurred in a few individuals or families across one or a couple of generations, they remain intrinsically limited in resolution due to the small number of recombination events that occur per chromosome per generation (Clark et al. 2010; Peñalba and Wolf 2020).

Another type of approach uses genome sequence data from population samples to take advantage of the large number of recombination events that have occurred during the history of the considered population. Instead of directly observing crossover products, these methods detect the footprints left by historical recombination events on patterns of haplotype segregation and linkage disequilibrium (LD) (reviewed in Stumpf and McVean 2003). The recombination rate and its variation across the genome are inferred via coalescent-based analysis of DNA sequence polymorphism data (Chan et al. 2012; Kamm et al. 2016; Li and Stephens 2003.; McVean et al. 2004; Spence and Song 2019). The resulting LD maps have been widely used to evaluate the genomic impact of natural selection and admixture, and to perform genome-wide association studies (GWAS) (*e*.*g*. Chan et al. 2012; The International HapMap Consortium 2007). These approaches provide an accessible and attractive way of describing recombination landscapes - i.*e*. the variation of recombination rates along the genome - particularly in non-model taxa where direct methods are often difficult to implement (Auton et al. 2012; 2013; Melamed-Bessudo et al. 2016; Shanfelter et al. 2019; Singhal et al. 2015; Shield et al., 2020).

Direct and indirect methods have revealed considerable variation in recombination rate at different scales along the genome, particularly in vertebrates. At a large scale (of the megabase order), recombination tends to be concentrated in subtelomeric regions compared to centromeric and centro-chromosomal regions, a pattern shared among many species of plants and animals (Auton et al. 2012; Melamed-Bessudo et al. 2016; Capilla et al. 2016; Danguy des Déserts et al. 2021; Haenel et al. 2018). At a finer scale (of the kilobase order), recombination events often cluster within small regions of about 2 kb, called recombination hotspots (Choi and Henderson 2015; Kim et al. 2007; Mancera et al. 2008; Myers et al. 2005; Singhal et al. 2015; Shanfelter et al. 2019; Schield et al., 2020). Two distinct regulatory systems of recombination hotspot location have been described to date, with major implications on the evolutionary dynamics of recombination landscapes. In passerine birds (Singhal et al. 2015), dogs (Axelsson et al. 2012; Auton et al. 2013) and some teleost fishes (Baker et al., 2017; Shanfelter et al., 2019), recombination hotspots tend to be found in CpG-islands / promoter-like regions, and are highly conserved between closely-related species (Singhal et al. 2015). In contrast, in humans (Myers et al. 2005; 2010), apes (Auton et al. 2012; Great Ape Genome Project 2016) and mice (Booker et al. 2017), hotspot location is directed by the PRDM9 protein, which binds specific DNA motifs and triggers DSBs (Oliver et al. 2009; Baudat et al. 2010; Myers et al. 2010; Parvanov et al. 2010; Grey et al. 2018). In these taxa, hotspots are mostly located away from genes (Auton et al., 2012; Baker et al. 2017), and show little or no conservation between closely related species (Myers et al. 2005, 2010; Auton et al. 2012; Booker et al. 2017) due to self-destruction by gene conversion and rapid turnover of PRDM9 alleles (Coop and Myers 2007; Lesecque et al. 2014; Latrille et al. 2017).

Population-based inference methods aim to infer the population recombination rate ρ = 4*N*_*e*_*r, r* being the per generation, per bp recombination rate and *N*_*e*_ the effective population size (Stumpf et McVean 2003). The *ρ* parameter reflects the density of population recombination events that segregate in polymorphism data, integrated across time and lineages. Several programs have been developed for reconstructing LD-maps (reviewed in (Peñalba and Wolf 2020; including PHASE: Li and Stephens 2003, LDhat: McVean et al. 2004, LDhelmet: Chan et al. 2012, LDpop: Kamm et al. 2016, and pyrho: Spence and Song 2019), which use the theory of coalescence with recombination to model the complex genealogies of samples stored in the underlying ancestral recombination graph (Griffiths et al. 1997; Arenas,2013). The most popular family of LD-based methods, comprising LDhat (McVean et al. 2004) and its improved version LDhelmet (Chan et al. 2012) (see a literature survey in Supplementary Figure S1), implement a pairwise composite likelihood method under a Bayesian framework using a reversible jump Markov Chain Monte Carlo (rjMCMC) algorithm. They have been used for building fine scale LD-based maps in a broad range of animal taxa including humans (McVean et al. 2004), dogs (Axselsson et al. 2012; Auton et al. 2013), fruit flies (Chan et al. 2012), finches (Singhal et al. 2015), honeybees (Wallberg et al. 2015), sticklebacks (Shanfelter et al. 2019) and rattlesnakes (Schield et al., 2020). In some species, inferred LD-based maps have been validated by comparison with recombination maps obtained using direct approaches, confirming their overall good quality (McVean et al. 2004; Chan et al. 2012; Singhal et al. 2015; Booker et al. 2017; Shanfelter et al. 2019). However, as genetic and LD-based maps greatly differ in their resolution (pedigree-based inference provide resolution of about 1 cM, while population-based methods can infer recombination events at the kilobase scale, Peñalba and Wolf 2020), such comparisons do not provide qualitative information on the reliability of inferred fine-scale variation and hotspots detection. Moreover, the heterogeneity of studies in terms of taxonomy, genetic diversity, demography, sample size, and software parameters, among other things makes it difficult to appreciate the performance and the possible weaknesses of LD-based methods. For these reasons, the reliability and conditions of application of LD-based methods are still poorly understood and need to be more thoroughly characterised, considering the growing importance of these tools.

The power and sensitivity of LDhat and LDhelmet have been tested by simulations aiming to evaluate the influence of switch error in haplotype phasing (Singhal et al. 2015; Booker et al. 2017), the amount of polymorphism, and the intensity of recombination hotspots (Singhal et al. 2015). These studies simulated simple recombination landscapes assuming either homogeneous recombination rates or a few, well-defined hotspots contrasting with a low-recombination background (McVean et al. 2004; Auton and McVean, 2007; Chan et al. 2012; Singhal et al. 2015; Booker et al. 2017; Shanfelter et al. 2019; Schield et al., 2020). Real recombination landscapes that were characterised with a fine-scale resolution such as in humans, however, appeared to be more complex and involve a continuous distribution of recombination hotspot density and intensity across genomic regions. This complexity has not been taken into account so far in benchmarking studies assessing the performance of LD-map reconstruction methods. We thus lack a comprehensive picture of the ability of these methods to properly recover the biological characteristics of human-like recombination landscapes interspersed with hotspots. In particular, the proportion of the inferred recombination hotspots that are correct, and the proportion of true hotpots that are missed, have not yet been quantified under a biologically realistic scenario. These are crucial quantities to properly interpret and use reconstructed LD-maps in genomic research.

In this work, we specifically assessed the performance of the LDhelmet program to detect hotspots while assuming a biologically realistic recombination landscape. We evaluated the influence of methodological parameters including sample size, phasing errors and block penalty, the impact of the population demographic history including its long-term effective size and the occurrence of bottleneck and admixture events, and finally the effect of the mutation to recombination rate ratio. We also considered different definitions of a recombination hotspot relative to its background recombination rate, with the aim of improving the sensitivity of the analysis. We identified the conditions in which LD-based inferences can provide an accurate mapping of hotspots, and the parameters that negatively affect the sensitivity and specificity of their detection across biologically realistic recombination landscapes.

## Results

### Recombination landscape modelling

Five realistic, heterogeneous recombination landscapes (referred to as “underlying landscapes” throughout) of 1Mb length were built using the human genome high resolution map of meiotic DSB from Pratto et al. (2014). In order to mimic both broad and fine-scale variation in the recombination rate parameter “*r*”, the first and second half of each landscape were drawn from a gamma distribution with mean 1 cM/Mb and 3 cM/Mb, respectively, and parameters fitted from Pratto et al. (2014) (1-500,000bp: shape=rate=0.1328; 500,001pb-1Mb: shape=0.1598, rate=0.0532). Accordingly, the 5 recombination landscapes generated (Supplementary Figure S2) showed broad-scale differences in recombination peak intensity, with less elevated recombination peaks in the first half compared with the second half of each chromosome. At a fine scale, recombination was concentrated in numerous peaks resembling human recombination hotspots, with about 85% of the recombination concentrated in 15% of the genome. The map lengths in recombination units were about 0.02 Morgan (Supplementary Figure S2, S4).

Population-scaled recombination landscapes simulated under a constant effective population size (hereafter called “simulated landscapes”) were generated in 10 replicates for the five underlying landscapes, using coalescent simulations with a mutation rate *μ*=10^-8 and 4 combinations of sample sizes (SS=10 or 20) and effective population sizes (*N*_*e*_=25,000 or 250,000) (Figure 1A, Supplementary Figure S3A). The map lengths of simulated landscapes were a little shorter than the underlying landscapes (about 0.015-0.018 Morgan), reflecting the occasional occurrence of more than one recombination event between two adjacent SNPs during the simulated coalescent histories (Supplementary Figure S4). These simulated landscapes were also highly correlated with the underlying landscapes for each combination of parameters (Spearman’s rank correlation > 0.8 using a 500bp resolution level), showing that the coalescent history has not resulted in a substantial loss of information about recombination rate variation across the underlying landscape. As expected from the *θ*=4*N*_*e*_*μ* values used in our simulations (*θ* = 0.001 and 0.01 for *N*_*e*_ = 25,000 and 250,000, respectively), the SNPs density of the large *N*_*e*_ populations was about one order of magnitude higher than for smaller *N*_*e*_ populations (Supplementary Figure S5).

**Figure 1.**
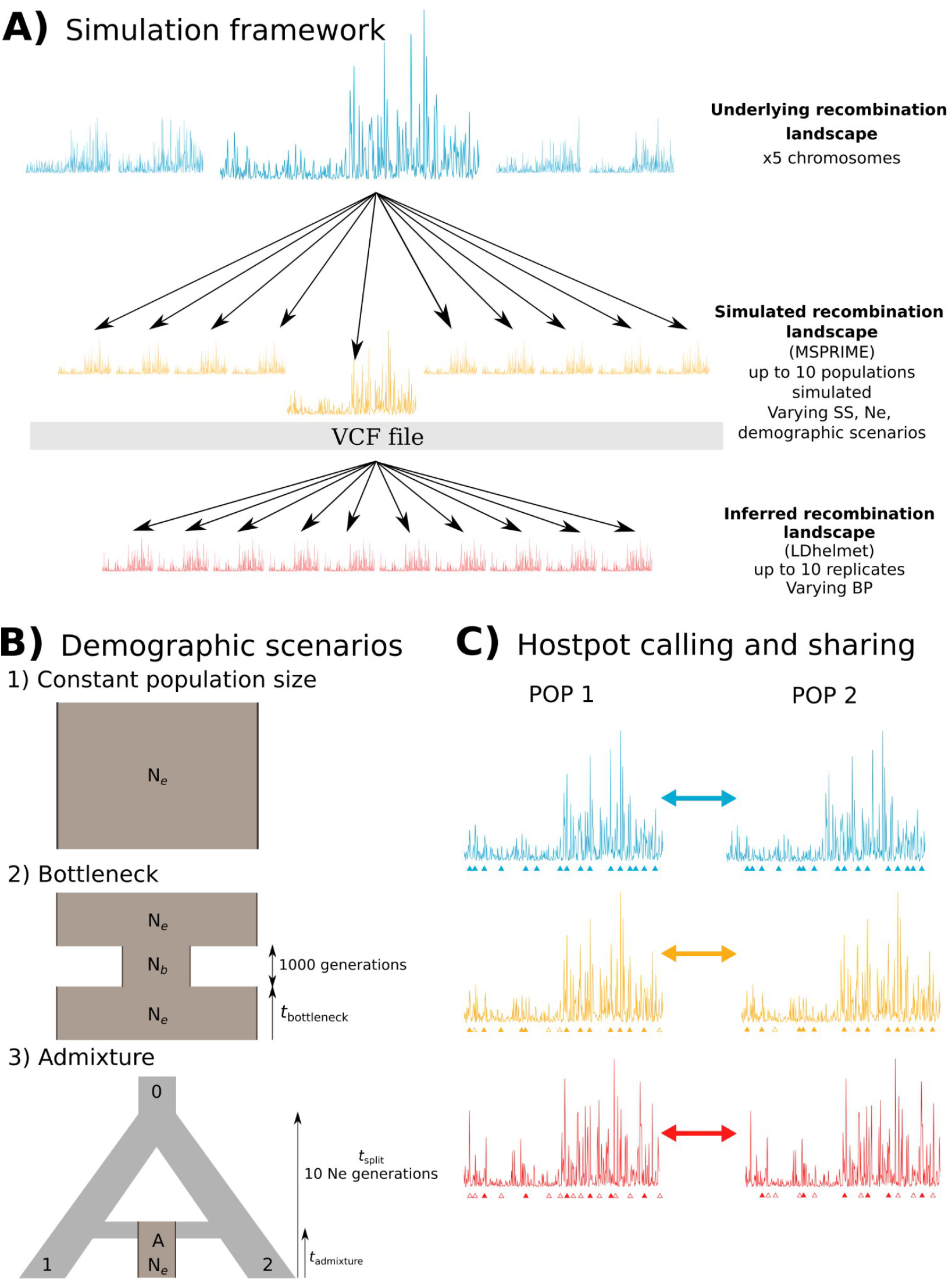
Simulation and hotspot calling protocols. **A)** Simulation framework. Five different underlying recombination landscapes were generated based on human empirical data. These five landscapes can either be considered as different regions from distinct chromosomes within a same species, or as orthologous regions of a same chromosome in different species. From these underlying landscapes, up to 10 recombination landscapes were simulated with MSPRIME 0.7.4 under various demographic scenarios, varying the effective population size (i.e. *N*_*e*_=25,000, 100000, 250,000) and the sample size (i.e. SS=10, 20), to generate a VCF file for each simulated population. The VCF files were then used to infer the local population recombination rates using LDhelmet 1.19 setting the block penalty to BP = 5, 10 or 50. Up to 10 replicates per simulated population were analysed with LDhelmet. **B)** Demographic scenarios. Three demographic scenarios were simulated with MSPRIME: 1) Constant population size, varying the *N*_*e*_ of the simulated population; 2) Bottleneck event, varying the age of the bottleneck (i.e. 500, 5000, 50000) and the *N*_*e*_ of the population during the bottleneck (i.e. N_b_=2500, 25,000), and setting the duration of the bottleneck to 1000 generations; 3) Admixture event, varying the age of the admixture event (i.e. 500, 5000, 50000), and setting the time of the split of the ancestral population into two populations 10*Ne generations ago. **C)** Hotspot calling and sharing. Hotspots in the underlying (blue), simulated (orange) and inferred (red) landscapes were defined as 2.5 kb-windows with a local recombination rate *X* times as high as the averaged recombination rate of the 50 kb flanking regions. Several threshold values were used to call the hotspots (i.e. *X*=2.5, 5, 10). The location of hotspots was compared between populations (that share or not the same underlying landscape), to compute the proportion of shared hotspots. Triangles below each landscape represent called hotspots, with filled triangles indicating shared hotspots, and empty triangles hotspots that are not shared with the compared landscape. In this example, the two compared populations share the same underlying landscape, meaning that all the hotspots are shared between them (all blue triangles are filled). The two simulated landscapes with independent coalescent histories share most of these hotspots, while the two landscapes inferred with LDhelmet share a smaller fraction of them, due to the additional inference step.

### Methodological parameters affecting LDhelmet performance

Population-scaled recombination rates (ρ) were inferred from the simulated polymorphism datasets using the program LDhelmet (Chan et al. 2012) (referred to as “inferred landscapes” throughout). The effect of sample size and landscape resolution level were assessed for a constant effective population size (*N*_*e*_=25,000) using 10 or 20 diploid individuals (SS=10 or 20) and three block penalty (BP) values (BP=5, 10 or 50), which inversely determine the number of allowed changes in ρ value within windows of 50 consecutive SNPs (Figure 1A, Supplementary Figure S3A). Underlying and simulated landscapes were converted into population-scaled recombination rates (ρ=4*N*_*e*_*r*), and each underlying, simulated and inferred maps was smoothed using 500bp (*i*.*e*. the underlying landscape resolution level) and 2500bp windows (*i*.*e*. a resolution level better suited to the SNP density in our low-*N*_*e*_ simulations). The 10 simulated and inferred replicates of each SS/BP condition were averaged to perform landscapes comparisons. Overall, local recombination rates tended to be overestimated by LDhelmet, no matter the value used for SS and BP, but this was especially observed when the local *ρ* was either very low (*ρ* < 10^-4) or very high (ρ > 10^-2) (Supplementary Figure S6, panels A-F). The mean inferred map lengths calculated across replicates varied substantially among tested conditions (0.017-0.125 M), reaching up to 6 times the length of simulated maps in low SS and BP conditions (*i*.*e*. SS = 10, BP = 5, 10, Supplementary Figure S4, upper panel). A BP value of 50 produced very smooth recombination maps, which did not capture fine-scale variation in recombination rate. By contrast, maps inferred with BP=5 or BP=10 were visually similar and better reflected the fine-scale variation of the underlying landscapes (Supplementary Figure S7A). Spearman rank correlation coefficient between the mean simulated and inferred landscapes was reduced when SS were small (Figure 2A). Replicate runs of LDhelmet showed a strong consistency, as revealed by elevated correlations among the 10 replicate landscapes inferred from the same simulated landscape, whatever the SS and BP values being tested (Spearman’s rho > 0.89, Figure 2B).

**Figure 2.**
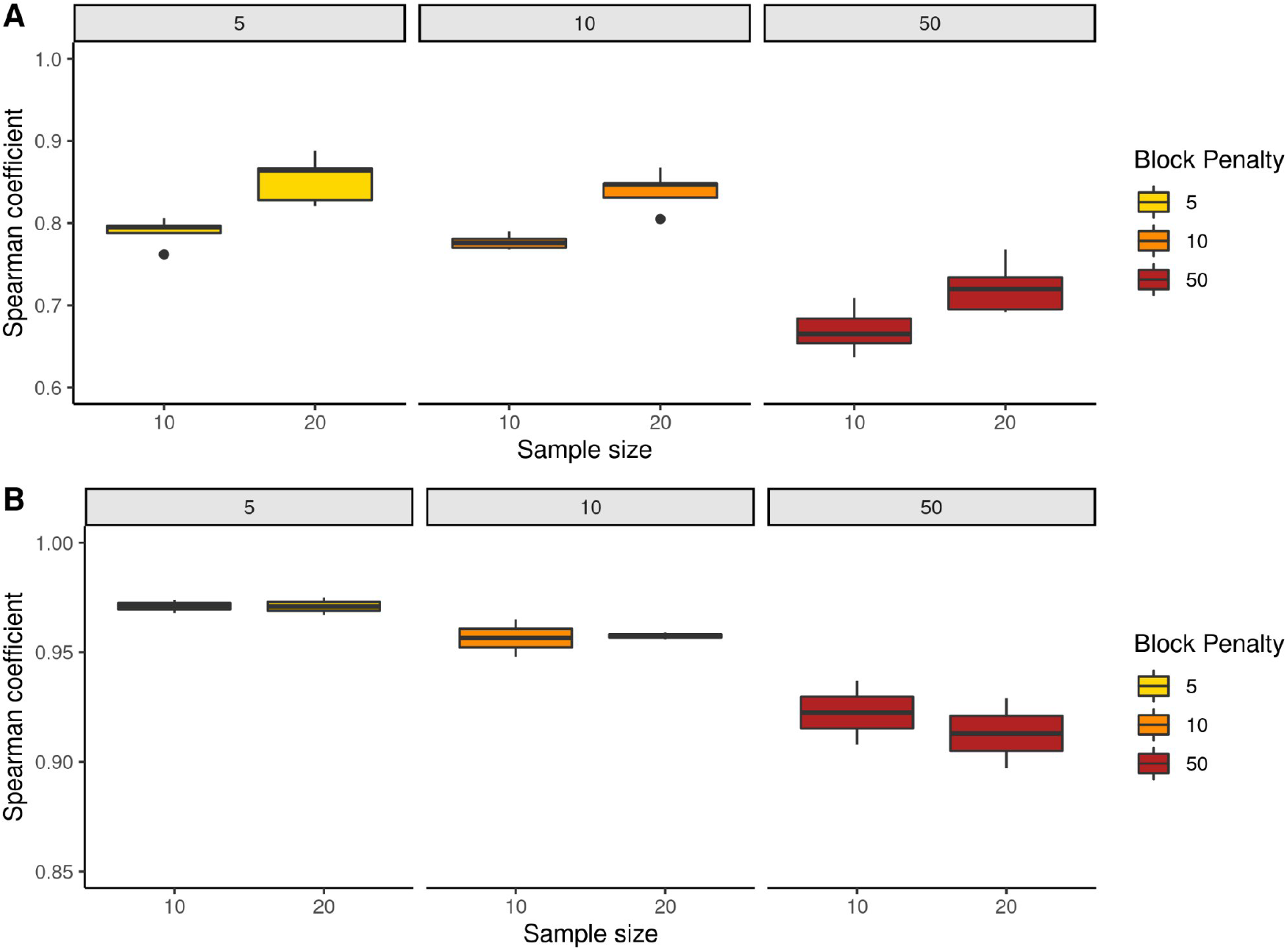
Performance (A), and repeatability (B) of LDhelmet as a function of the different parameters tested (i.e. *N*_*e*_, SS, BP). The *N*_*e*_ of the simulated population is 25,000, the sample size is shown on the x axis (i.e. SS=10 or 20), and the LDhelmet BP values shown in color correspond to the different panels (i.e. BP=5, 10 or 50). **A)** Spearman correlation coefficients between the mean simulated and the mean inferred landscape calculated across the 10 replicates obtained from each of the five underlying landscapes (i.e. using simulation framework of Supplementary Figure S3A). **B)** Mean pairwise Spearman correlation coefficients calculated between the 10 replicates of inferred landscapes from simulated populations sharing the same underlying landscape (i.e. using simulation framework of Supplementary Figure S3B).

Recombination hotspots of the underlying, simulated and inferred landscapes were called using three different threshold values commonly used in the literature (*i*.*e*. local recombination rate at least 2.5, 5 or 10 times higher than the background rate). True/False positives/negatives rates and discovery rates (TPR, FPR, TDR, FDR, TNR, FNR) were computed under each tested condition. The hotspot detection threshold ratio of 10 between the focal and flanking positions appeared too stringent and yielded a very small number of called hotpots (Supplementary Figure S8). Using a less conservative threshold ratio of 5, we detected 4 to 8 hotspots per Mb in the simulated landscapes, and 5 to 20 per Mb in the inferred landscapes. These numbers reached 40-50 and 20-50 per Mb, respectively, when a threshold of 2.5 was used. Irrespective of the chosen threshold, the number of inferred hotspots tended to be overestimated, notably when SS was small (Supplementary Figure S8). The 2.5 threshold was used for the remaining analysis as it reduced the variance in the number of called hotspots due to a higher call rate. The sensitivity (or TPR) of LDhelmet was medium, since depending on the SS and the BP used, between 29.4% and 52.7% of the simulated hotpots were inferred as such. The TPR was higher for small BP values (*i*.*e*. 5 or 10), but relatively insensitive to the SS value (Figure 3 left panel, Supplementary Figure S9A). The proportion of false hotspot calls (FDR, *i*.*e*. inferred hotspots corresponding to non-hotspot windows in the simulated maps) ranged between 25.6% and 52.9%, and was higher for SS = 10, without major differences between the BP values tested (Figure 3 right panel, Supplementary Figure S9B and C). No significant difference in the correlation between simulated and inferred landscapes was found between the first half of the chromosome with a mean r of 1 cM/Mb (referred to as the “cold” region) and the last half with a mean *r* of 3 cM/Mb (the “hot” region). This was also true for the TPR and the FDR, whatever the hotspot detection threshold used (*i*.*e*. 2.5 or 5) (Student test, p > 0.05).

**Figure 3.**
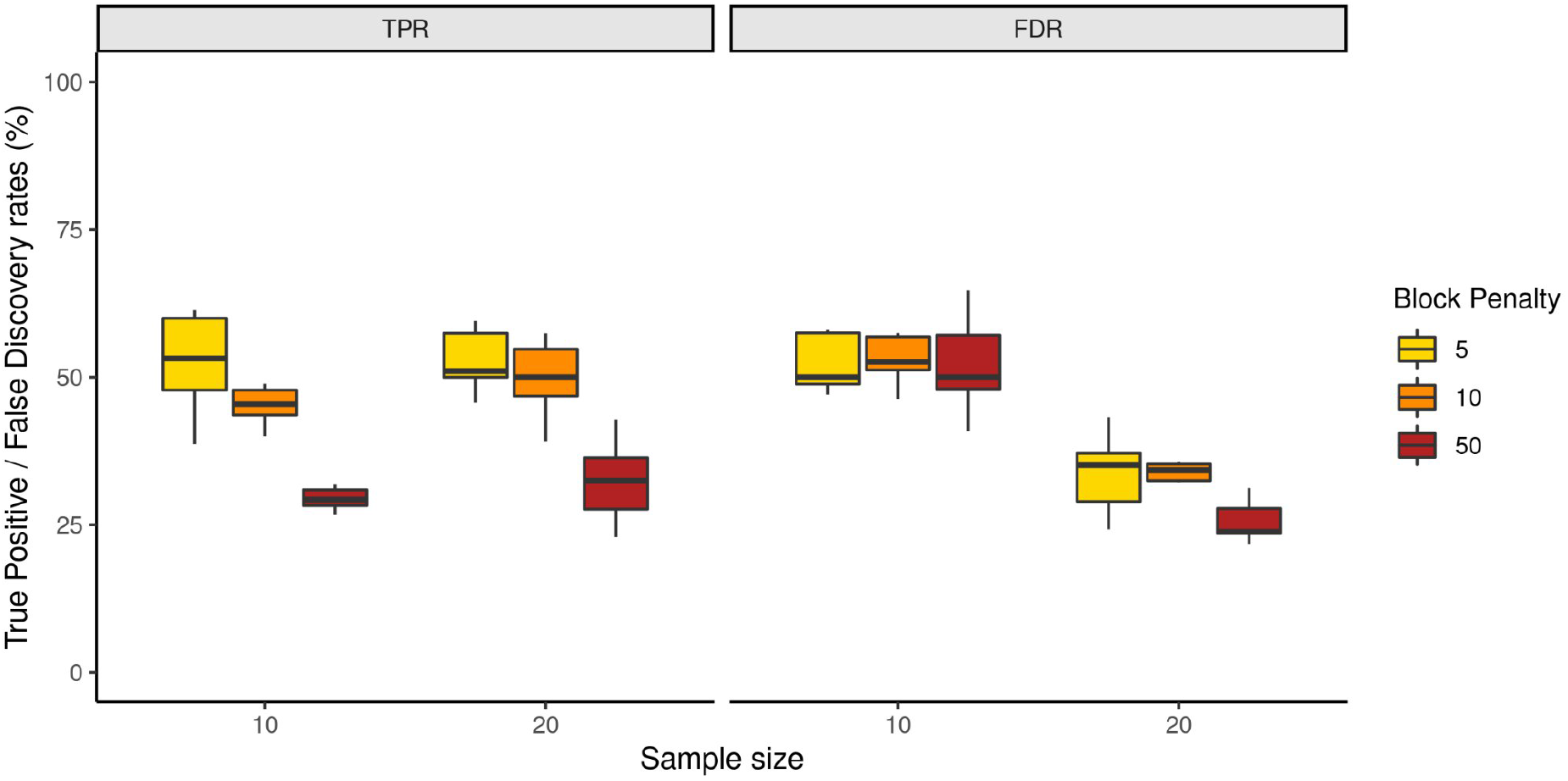
Hotspot detection. True positive (TPR, left panel) and false discovery (FDR, right panel) rates of inferred, as compared to simulated hotspots, called using a detection threshold of 2.5, for different sets of methodological parameters. The sample size parameter is shown on the x axis (i.e. 10 or 20), the block penalty (i.e. 5, 10, 50) is shown in color, and the *N*_*e*_ of the populations simulated is 25,000. See full results in Supplementary Figure S9.

To assess the impact of phasing errors on LDhelmet performance, the phase information was first removed from the whole VCF of the simulated landscapes for two replicate populations simulated with a constant Ne of 250,000 and a SS of 20. Statistical phasing performed with Shapeit 4.2.2 (Delaneau et al. 2019) resulted in a 6.7% average phasing error. We then introduced random phasing errors in the simulated VCFs with rates ranging from 2 to 10%, before inferring ρ with a BP value of 5. As the phasing error rate increased, Spearman rank correlation coefficients between simulated and inferred landscapes slightly decreased below the value obtained with perfectly phased data (Supplementary Figure S10A). Hotspot calling performance assessed with TPR and FDR was also negatively impacted by phasing errors (Supplementary Figure S10B).

### Demographic and evolutionary parameters affecting LDhelmet performance

Methodological parameters were then set to SS=20 individuals and BP=5 - a trade-off optimising the balance between TPR and FDR - to focus on the effect of *N*_*e*_ on the quality of the LD inferences. When the simulated effective population size was large (i.e. 250,000, as compared to 25,000), the inferred map length was closer to the expected value of 0.02 M (Supplementary Figure S4, lower panels) and the local recombination rate tended to be less overestimated (Supplementary Figure S6, panels G-L). A larger *N*_*e*_ also significantly increased the correlation between simulated and inferred landscapes (Figure 4A, Supplementary Figure S11), increased the TPR and decreased the FDR (Figure 4B, C).

**Figure 4.**
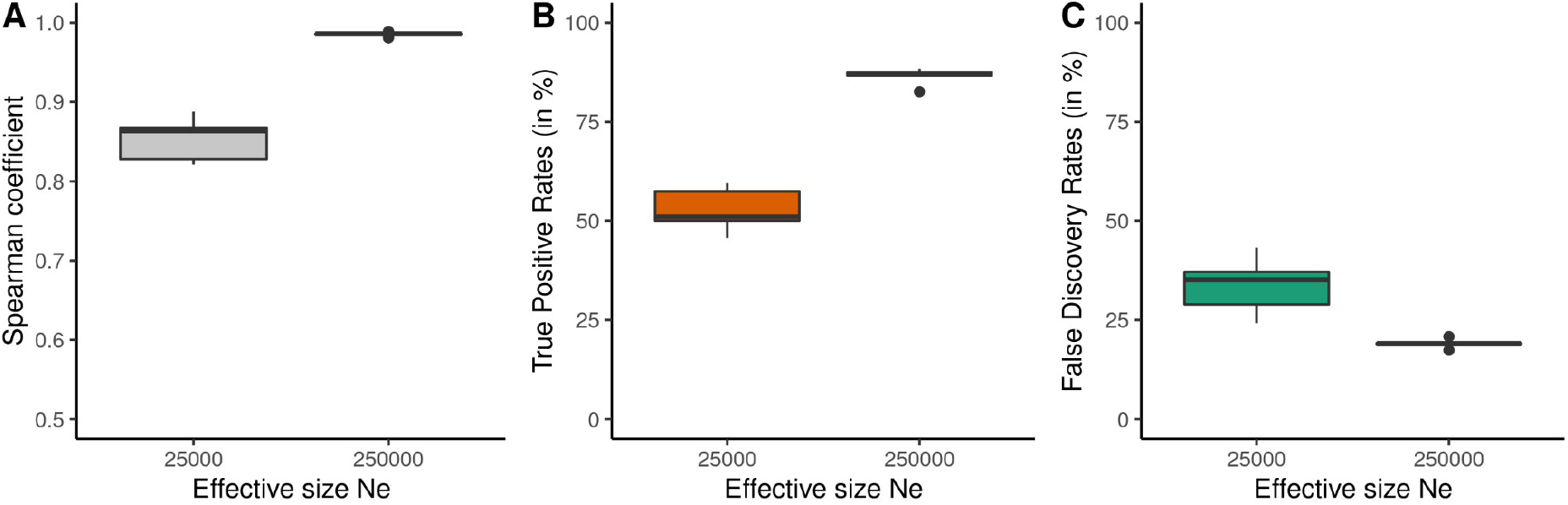
Effect of the effective population size parameter (Ne) on recombination rate inference and hotspot detection. **A)** Spearman correlation coefficients between simulated and inferred landscapes as a function of *N*_*e*_ (SS = 20, BP = 5). **B)** and **C)** True positive (TP) and false discovery (FD) rates of inferred, as compared to simulated hotspots, called using a detection threshold of 2.5, as a function of *N*_*e*_ (SS = 20, BP = 5). See full results in Supplementary Figure S9 and S11.

We then assessed the impact of non-equilibrium demographic histories. Populations undergoing bottleneck or admixture events of various ages were simulated, with the *N*_*e*_ of the ancestral and present-day populations set to 250,000, the SS to 20, and making other demographic parameters vary (*i*.*e. t*_b_, *N*_b_ *t*_a_, Figure 1B, see Materials and Methods). These demographic scenarios generally had a negative impact on the quality of the reconstructions (using a BP of 5), as compared to constant-size populations (Figure 5). The correlation between simulated and inferred maps decreased with the strength of the bottleneck (i.e. with lower *N*_*b*_), and with the recentness of bottleneck and admixture events (Figure 4A, B). In the same way, TPR and FDR were degraded as compared to constant-size population scenarios (Figure 4C, D), particularly for young events.

**Figure 5.**
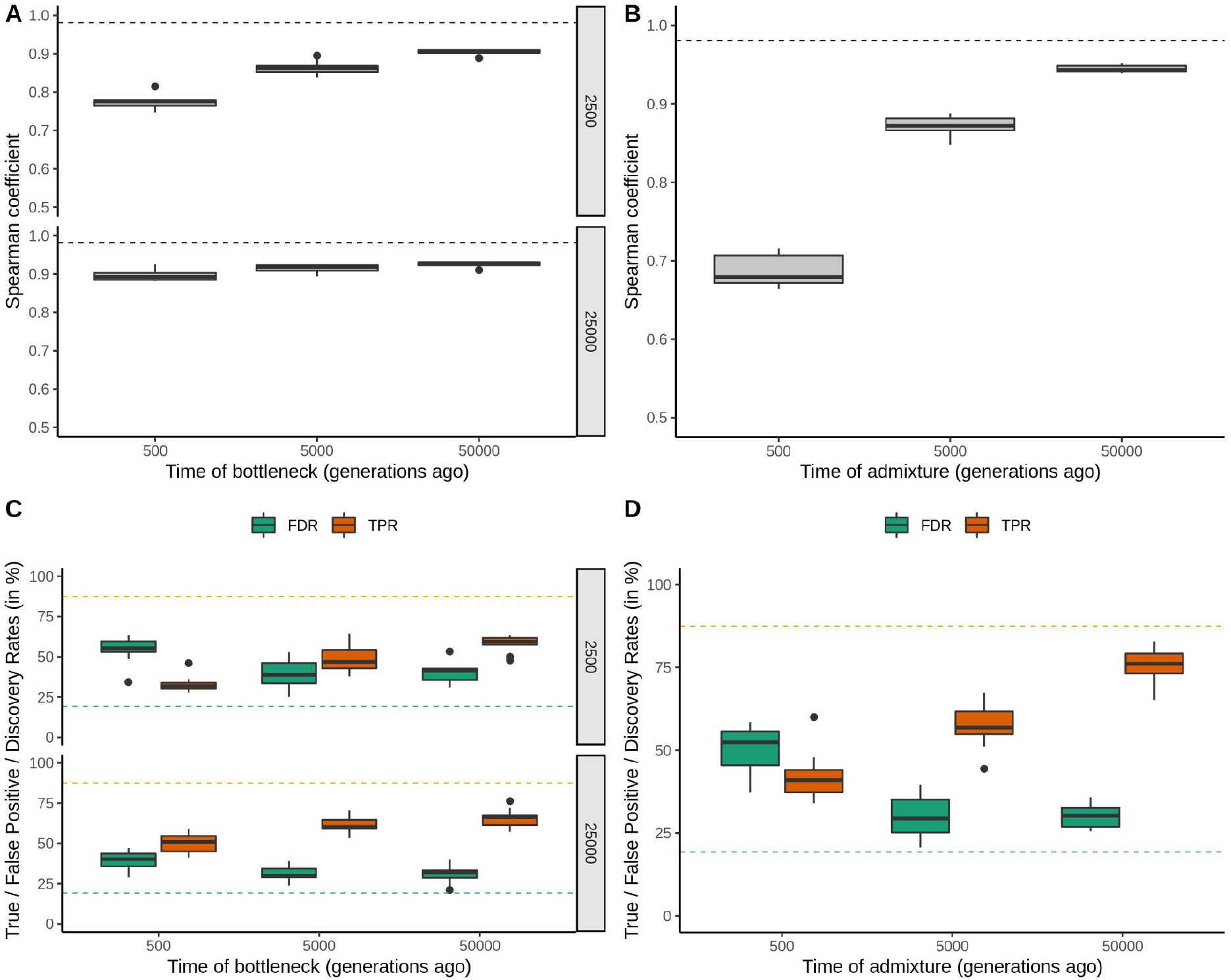
Influence of bottleneck (left panels A and C) and admixture (right panels B and D) events on recombination rate inference and hotspot detection. Spearman correlation coefficients between simulated and inferred landscapes are shown as a function of the age of the bottleneck event and the strength of the bottleneck (i.e. *N*_*b*_, the *N*_*e*_ value during the 1000 generations of the bottleneck) (**A**), and as a function of the age of the admixture event (**B**). True positive (TP, in orange) and false discovery (FD, in green) rates of hotspots called using a detection threshold of 2.5, as a function of the time of the bottleneck event and the strength of the bottleneck (**C**), and as a function of the time of the admixture event (**D**). Dashed lines correspond to averaged Spearman’s rho, TPR and FDR values in populations that did not experience bottleneck or admixture events.

The influence of species-specific evolutionary parameters such as the mutation and recombination rates was assessed by generating coalescent simulations under two additional underlying landscapes using a ten times higher (*i*.*e*. 20 cM/Mb) and a ten times lower (*i*.*e*. 0.2 cM/Mb) average recombination rate, and three different mutation rates (*i*.*e*. 10^-9, 10^-8 and 10^-7). The *μ*/*r* ratio under these 6 simulated conditions thus equalled 0.1, 1 or 10. For all conditions, Spearman’s rank correlation between the mean simulated and the mean inferred landscapes was greater than 0.9, except when *μ* equalled 10^-9 (Spearman’s rho ≃ 0.7, Supplementary Table S1). An increased *μ*/*r* ratio improved the ability to detect hotspots when *r* was fixed to 10^-8, with a higher TPR (up to >80%) and a lower FDR (<5%) when *μ* increased (prop.test, p-value < 0.05, Figure 6, threshold 5 in Supplementary Figure S12). The *μ*/*r* ratio did not affect the performances the same way when *μ* was fixed to 10^-8: a *μ*/*r* ratio of 10 (r = 10^-9) yielded lower TPR (< 60%) and higher FDR (> 25%) than a ratio of 1 or 0.1, although these trends were not significant (prop.test, p-value > 0.05, Figure 6, threshold 5 in Supplementary Figure S12).

**Figure 6.**
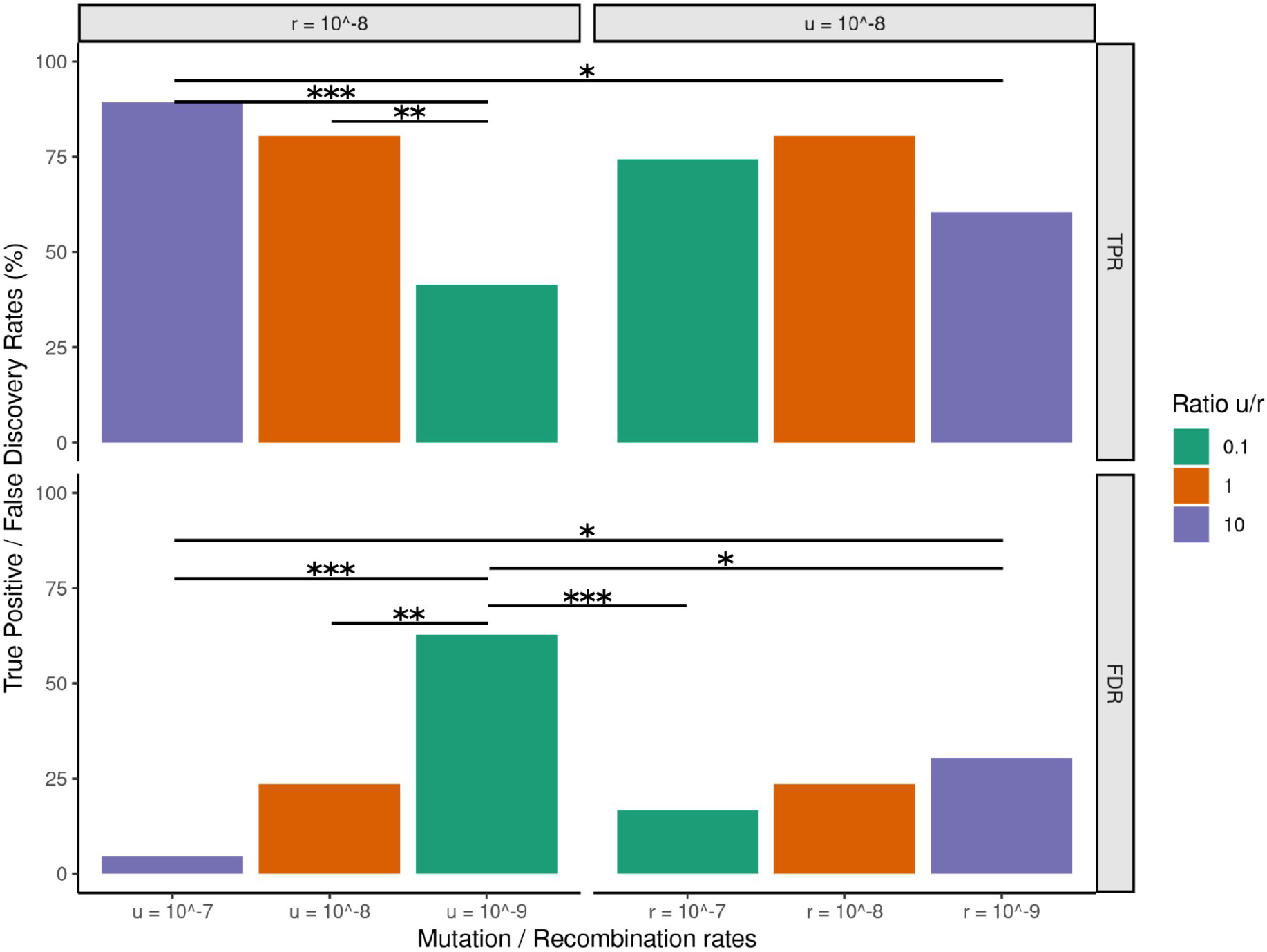
Influence of the u/r ratio on hotspot detection. True positive (TP, upper panel) and false discovery (FD, lower panel) rates of hotspots called using a detection threshold of 2.5. The x axis indicates values of u for *r*=10^-8 (left panels), and values of *r* for *μ*=10^-8 (right panels). Colors correspond to different values of the *μ*/*r* ratio. Asterisks show significant differences in percentages between comparisons (* prop.test p-value<0.05, **<0.01, ***<0.001).

### Hotspot sharing between populations with different versus identical underlying recombination landscapes

As expected for a comparison between two populations simulated with different underlying recombination landscapes, the mean linear correlation (R^2^ coefficient) between the corresponding inferred landscapes was low, between 0.012 and 0.084, and similar to the R^2^ between the simulated landscapes (0.012-0.017) (Supplementary Table S2). A low percentage of shared hotspots (around 8% with a calling threshold of 2.5) occurred by chance between populations simulated with distinct underlying landscapes, with a SS of 20. Roughly similar proportions of shared hotspots were found between the corresponding inferred landscapes (with BP = 5), although these proportions were slightly overestimated (Figure 7, see Supplementary Table S2 to see all conditions). A minority of the shared inferred hotspots were TP, indicating that a non-zero fraction of truly shared hotspots is expected to be found between species with different biological recombination landscapes.

**Figure 7.**
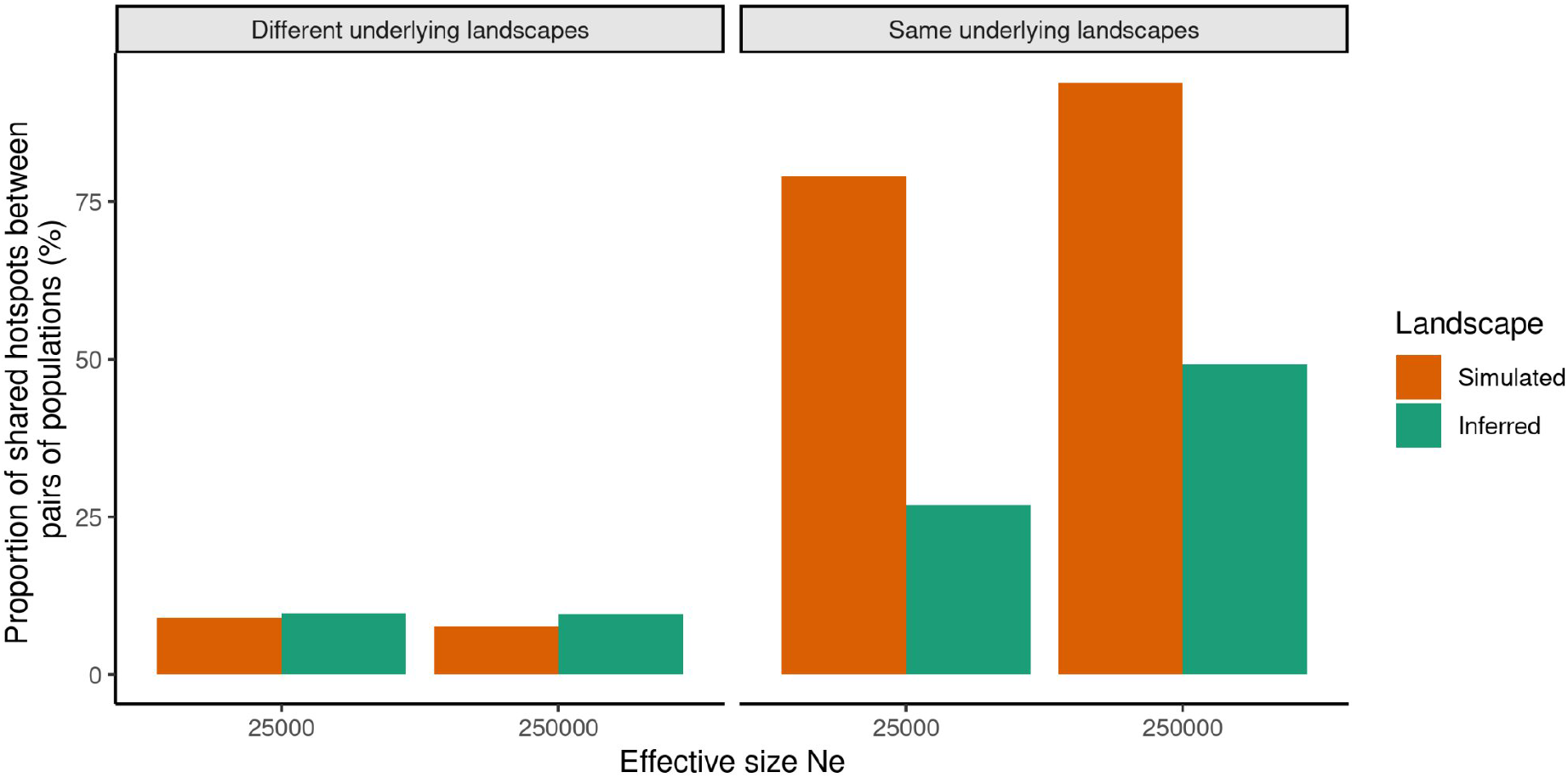
Expected and observed hotspot sharing between populations with different (left panel, following simulation framework from Supplementary Figure S3 A)) or identical (right panel, following simulation framework from Supplementary Figure S3 B) underlying landscapes. Mean proportion of shared hotspots between pairs of simulated (expected proportion, orange bars) and pairs of inferred (observed proportion, green bars) recombination landscapes as a function of *N*_*e*_ (i.e. 25,000 and 250,000, x axis). Only shown here are simulations with SS = 20, BP = 5, hotspots detection threshold = 2.5. The proportion of shared simulated and inferred hotspots for all combinations of parameters and for populations sharing or not the same underlying landscape are shown in Supplementary Table S2.

Then we compared simulated populations sharing the same underlying landscape, in order to check the ability of LDhelmet to recover similar recombination rates between populations with independent coalescence histories. The correlations between simulated landscapes were generally high for both low (R^2^>0.7) and large *N*_*e*_ (R^2^>0.9) conditions, but the correlations between inferred landscapes were much lower, with R^2^< 0.3 and <0.75 for *N*_*e*_ = 25,000 and 250,000, respectively (Supplementary Table S2). The proportion of shared hotspots called with a threshold of 2.5 followed the same trend: it was high between simulated landscapes (>80%) and much lower between inferred landscapes (<50%) (Figure 7, Supplementary Table S2). Thus, one can expect LDhelmet to detect a moderate to low fraction of shared hotpots even between species truly sharing a common recombination landscape, depending on population size and sample size (and also hotpot definition).

Populations simulated with the same underlying landscape that underwent a bottleneck or an admixture event also showed reduced hotspot sharing between inferred landscapes, as compared to truly shared hotspots between simulated landscapes. The proportion of shared hotspots between pairs of simulated landscapes was similar to constant-size populations and did not vary substantially according to the time or the strength of the demographic event, while the proportion of shared inferred hotspots decreased for younger events. This proportion was overall lower than for constant-size populations, but it sometimes reached, or even exceeded, the constant-*N*_*e*_ reference when events were ancient (Supplementary Figure S13 G, H).

## Discussion

### Inferred LD-maps should be interpreted with caution

Inference methods based on linkage disequilibrium provide an attractive way to characterise genomic recombination landscapes from sequence data. As such, they promise to become increasingly popular in empirical studies of eukaryotes. However, their ability to accurately reproduce real recombination landscapes has not been specifically evaluated. It should be recalled, however, that LD-based recombination maps are inferences, not observations; biases and uncertainty must be quantified and taken into account when it comes to interpreting the results. Here, we modelled the biological characteristics of a particularly well-documented recombination landscape, that of humans, as a basis for assessing the impact of methodological and species-specific demographic and evolutionary parameters on the performance of the LDhelmet method. Our results send a message of caution regarding the reliability of reconstructed recombination maps and hotspot location.

Indeed, we show that the recombination landscapes inferred with LDhelmet differ from real landscapes, sometimes substantially, with Spearman rank correlation between simulated and inferred 2.5 kb windows sometimes getting as low as 0.7 (Figure 2A, Supplementary Figure S11A). Hotspot detection is a particularly tricky and error-prone task: while up to 85% of true hotspots can be detected in the most favourable situations (*N*_*e*_ = 250,000, SS = 20, BP = 5, threshold = 2.5), the FDR ranged from 19% to 82% (Figure 3, Figure 4B and C, Supplementary Figure S9) according to the type of data and parameters used, meaning that in many cases a majority of the detected hotspots are incorrect calls. These discrepancies lead to a reduction in the apparent overlap in hotspot location between species/populations with identical recombination landscapes, while in turn inflating apparent hotspot sharing in populations with divergent landscapes (Figure 7, supplementary Table S2). These results were obtained with recombination landscapes of simulated populations with constant effective sizes, and with perfectly phased data. In reality, empirical data needs to be phased, and the phasing process can be prone to errors. Our analyses suggest that the typical error rate of statistical phasing methods such as Shapeit4 is relatively low (∼ 6.7% in our simulations) and only marginally affects the performance of LDhelmet in terms of sensitivity and hotspot sharing (Supplementary Figure S10), which is quite reassuring. Moreover, studied populations often hide complex demographic histories that are known to impact the power to correctly infer recombination rates (Dapper and Payseur 2018, Samuk and Nook 2021). We showed that recent bottleneck and admixture events tend to decorrelate simulated and inferred landscapes, decrease the TPR and increase the FDR, thus increasing the difficulty to call shared hotspots between populations sharing the same underlying landscape (Figure 5, Supplementary Figure S13). The significant impact of non-equilibrium demographic histories illustrated by our simulations provides additional motivation to characterise these histories in comparative studies of recombination landscapes. These results also provide a qualitative assessment of the impact of linked selection, whose effects may be similar to a local reduction in *N*_*e*_ (i.e. purifying selection), or to the maintenance of anciently diverged alleles (i.e. balanced selection, possibly involving structural variants such as chromosome inversions). If neglected, these effects might mislead biological interpretations regarding the evolutionary conservation of recombination maps.

In a study of the short time-scale dynamics of recombination landscapes based on LDhelmet, Shanfelter et al. (2019) found only 15% of shared hotspots between two recently-diverged populations of threespine stickleback. A greater overlap in hotspot location was *a priori* expected given that this species lacks a functional PRDM9 protein, which is responsible for the rapid turnover of recombination landscapes in mammals (Axelsson et al. 2012; Paigen and Petkov 2018). The authors suggested that a new mechanism of recombination hotspot regulation, different from the two already described in the literature, might be at play in this teleost species. In the light of our results, however, one cannot exclude that the strong divergence between the two reconstructed landscapes is due to a lack of power of the method in the first place. While the sample size of both fish populations was at least 20 individuals, *θ* was about 0.002, similar to our simulated conditions with a low *N*_*e*_. Under these conditions, a high FDR and a low proportion of shared hotspots can be expected even if the true underlying maps are identical (Supplementary Figure S9, Supplementary Table S2).

It should be recalled that real data sets typically carry less signal and more noise than simulated data sets, meaning that our assessment of the reliability of LDhelmet might be an overoptimistic one. In particular, our data sets are immune from sequencing errors or mapping errors, all of which presumably make the problem of recombination map inference an even harder one.

### Guidelines for population-based inference of recombination maps

Our study revealed that whatever the parameters used, the inference of recombination rates by LDhelmet is more reliable for species with large as compared to small effective population size (Figure 4, 7, Supplementary Figure S9, S11). This might be expected since long-term *N*_*e*_ determines the amount of nucleotide diversity (θ=4*N*_*e*_*μ*, Watterson 1975), so that a higher *N*_*e*_ results in a higher SNP density and a finer scale characterization of the recombination rate variation along the genome. Moreover, a higher effective size greatly corrects the general tendency of LDhelmet to overestimate the ρ value, especially for low and high recombination rates (Supplementary Figure S6, Singhal et al. 2015; Booker et al. 2017). Thus, when studying species with heterogeneous effective population sizes in nature, it is recommended to select populations with the largest *N*_*e*_, for which genetic diversity is greater. The question is then: how to obtain a good-quality recombination map when dealing with low *N*_*e*_ species? The sampling effort also determines, to a lesser extent, the polymorphism level of the dataset (Supplementary Figure S5), improving the accuracy of the inference (Figure 2, 3 Supplementary Figure S9, S11). A sample size of 20 is recommended based on our simulations. Moreover, as previously mentioned (Chan et al. 2012; Singhal et al. 2015), the block penalty parameter of LDhelmet, which determines the resolution level of the inferred landscape, also influences the length of the inferred map (*i*.*e*. a higher BP tends to mitigate the tendency of LDhelmet to overestimate the map length) and the number of detected hotspots (Supplementary Figure S4, S8). A small BP, that allows more fine-scale changes in the inferred ρ value, should be used to detect recombination hotspots. The ability of LDhelmet to faithfully reflect the fine-scale variation of real recombination landscapes is of great importance when it comes to detecting recombination hotspots. To this purpose, the threshold used to decide which region is defined as a “hotspot” is a key parameter that determines the level of detection stringency. If the chosen value is not appropriate, LDhelmet will detect false positives while also missing true hotspots (Supplementary Figure S9). This threshold should thus be adapted to the species studied, using a less stringent threshold in species with lower genome-wide average recombination rate.

Other intrinsic biological variables influence the ability to produce a faithful recombination map, such as the *μ*/*r* ratio, which in part determines the power to measure ρ at a fine-scale. The among-species variations in genome-average recombination rate *r* is well documented, ranging from 0.01 to 100 cM/Mb in animals and plants, with vertebrate taxa displaying an average *r* around 1 cM/Mb (Stapley et al. 2017). As previously mentioned, high and low recombination rates tend to be overestimated by LDhelmet, thus the average *r* of the studied species is obviously a key parameter to account for. The mutation rate *μ* also has a key impact on the performance of LDhelmet, since ancestral recombination events can only be detected if properly tagged by flanking mutations. The variation in *μ* across taxa, and consequently the ratio of *μ*/*r*, are much less well known than the variation in *r*. This ratio, which does not depend on the effective size of the population, is about 1 in humans, which means that two recombination events are separated by one mutation on average. A ratio in favour of mutations (*μ*/*r* > 1) will improve the signal, increasing the TP rate and reducing the FD rate (Figure 6, Supplementary Figure S12). But ultimately the performance of LDhelmet is conditioned by *r*, as low *r* values provide less power to detect the recombination events, even with *μ*/*r* = 10. Thus, the mutation to recombination rate ratio is crucial to build a non-biased recombination map. When studying a species for which it appears that this ratio is not favourable, a high rate of false positive hotspots is expected from the inferred population recombination landscape (Figure 6), making it difficult to compare maps between closely related species in a meaningful way.

### Limitations

The aim of our study was to determine the limits of LD-based methods in inferring biologically realistic recombination landscapes. For this purpose, we used the Pratto et al. (2014) ChIP-seq DMC1 data set to build human-like recombination landscapes including both broad and fine scale variation, reflected by the presence of numerous recombination hotspots of different intensities (Supplementary Figure S2, Myers et al. 2005, 2006; Pratto et al. 2014). We therefore assumed that the distribution of DSB reflects the distribution of crossing overs, which is not true for sure. For instance, hotspots were here placed without taking into account the existence of genomic features that correlate with the recombination rate, such as genes and promoter-like regions, GC-rich regions, CpG islands, and polymorphic regions, which can explain why a very intense and narrow hotspot is never found within a region of near zero recombination. The sensibility of LD-based methods with respect to this architecture was not tested. We did not take into account the effect of gene conversion on the dissipation of LD in high-recombining regions. While recent methods aim to distinguish between crossing-over (CO) and non-crossing-over (NCO) events (heRho, Setter et al. 2022), they do not (yet) account for the small-scale heterogeneity of recombination rates, and so are not really applicable when it comes to differentiate hotspots and NCO. Our simulated data were perfectly polarised, without missing or low-quality genotypes, which can’t be the case when dealing with empirical data. We simulated phasing errors in order to assess the robustness of LDHelmet to this problem. However, we estimated the phasing error rate of Shapeit4 from our simulated data which lack most of the biases found in empirical data, thus probably underestimating the typical phasing error of this method.

Finally, we don’t know if these simulated landscapes are representative of the diversity of recombination landscapes that exist in the living world (*i*.*e*. PRDM9-dependent vs independent landscapes, hotspot-free landscapes…). Indeed, it is likely that the high complexity of the human recombination landscape is not a universal feature in the animal kingdom. Singhal et al. (2015) used LDhelmet for building the recombination map in two species of birds, the zebra finch and the long-tailed finch, that lack a full-length PRDM9 gene copy and diverged about 2.9 Myr. The sample size for both populations was about 20 individuals, and θ (∼ 0.01) was about ten times higher than in apes or the threespine stickleback (Shanfelter et al. 2019), thus corresponding to our high *N*_*e*_ simulation conditions. Singhal et al. (2015) found 73% of shared hotspots between the two finch species, which is a higher rate of hotspot sharing than in any of the scenarios we simulated. The median estimated recombination rate was 0.14 cM/Mb in both species of finch, which is seven times lower than the genome-wide average recombination rate in humans (about 1 cM/Mb, Jensen-Seaman et al. 2004). Combined with the strong polymorphism in those species, we may suppose that birds possess less complex recombination landscapes than humans or compared to what we simulated, which might explain why LDhelmet recovered such a high percentage of shared hotspots in this study.

## Conclusion

In the past few years, we have seen a growing interest in recombination rate estimation in functional and evolutionary genomics. Indirect, LD-based approaches raise methodological challenges that are addressed by sophisticated methods such as LDhat or LDhelmet, the reliability of which is still poorly characterised. Our study provides guidance to users of these methods based on the characteristics of their species, and calls for caution when it comes to interpreting fine-scale differences in recombination rates between species. Extending this approach to a more diverse set of underlying recombination landscapes would help characterise further the reliability of these methods and their range of applicability across data sets and taxa.

## Materials and Methods

Our approach separately considers three different layers of information that are involved in the study of recombination landscapes (Figure 1). The first layer that we call the “underlying” recombination landscape corresponds to the true biological distribution of recombination rate (*r*) across the considered genome. We here used experimental measurements from human studies to model and generate the “underlying” landscapes. The second layer, the population recombination landscape, describes the genomic location of recombination events that occurred during the history of the sample. We used coalescent simulations to produce these population recombination landscapes, thereafter called “simulated” landscapes. Simulated landscapes differ from the underlying landscape due to the stochasticity of the coalescent process, which is inversely proportional to *N*_*e*_. The third layer, called the “inferred” landscape, corresponds to the output of LDhelmet, *i*.*e*. an estimate of the population recombination rate between adjacent SNPs. In total we generated five independent replicates of underlying landscapes, and for each of them up to 10 simulated and inferred landscapes under various demographic scenarios (Figure 1, Supplementary Figure S3A).

### Underlying landscapes

Underlying recombination landscapes were first generated to reproduce the features of the human recombination landscape. These include large-scale variation in the mean background recombination rate and fine-scale variation reflecting the presence of hotspots with varied intensities. Meiotic DSB are the major determinant of crossing over (CO) location along the genome (Li et al. 2019; Pratto et al. 2014). We used the high-resolution map of meiotic DSB obtained using ChIP-seq DMC1 in 5 non-related human genomes (Pratto et al. 2014) to define the genome-wide distribution of recombination rates in our simulations. The five individuals analysed in Pratto et al. (2014) carried different PRDM9 genotypes totalizing about 40,000 hotspots per individual, with distinct genotypes having different sets of DSB hotspots. For each individual, a gamma distribution was fitted to the empirical distribution of hotspot intensity measured by ChIP-seq DMC1 with the R package *figdistribplus* (Delignette-Muller and Dutang 2015). Extreme ChIP-Seq intensity values (>500) lying above the 97.5th quantile and likely representing technical artefacts were removed. Remaining values were rescaled to 0-100, so as to transform ChIP-Seq intensity values into quantities reflecting the range of recombination rates reported in cM/Mb across the human genome (McVean et al. 2004, 2005). This conversion assumed a linear relation between DMC1 activity and CO frequency (Pratto et al. 2014). We then removed null values and replaced them with small but non-null values (0.001), so that the genome-wide mean recombination rate equaled a target average (*e*.*g*. 1 cM/Mb). A Gamma distribution was fitted to these transformed empirical values separately for each of the 5 individuals, before averaging shape and scale parameters across individuals. Targeted genome-wide average value was set to either 1 cM/Mb or 3 cM/Mb, respectively reflecting the average centro-chromosomal and subtelomeric rates in humans. Underlying landscapes of 1 Mb length were built by randomly drawing independent recombination rate values from the fitted distribution and assigning these to non-overlapping windows of 500 pb. Values in the first 500 kb were drawn from a distribution of mean 1 cM/Mb, while values in the last 500 kb were drawn from a distribution of mean 3 cM/Mb. Our approach thus mimics both the large scale variation in recombination rate existing in humans (Nachman, 2002; Myers et al. 2005; Buard and de Massy 2007; Pratto et al. 2014) and the nearly absence of recombination events outside hotspots (96% of CO occur in hotspots in mice, Pratto et al. 2014; Li et al. 2019). In total, 5 underlying landscapes were generated (mean *r* = 2 cM/Mb), which can be considered as independent replicates driven from the same distribution (*i*.*e*. regions from different chromosomes of the same species, or orthologous chromosome region from closely related species).

### Simulated landscapes

For each of the 5 underlying landscapes, 10 simulated landscapes were generated via coalescent simulations using the program MSPRIME (v0.7.4, Kelleher et al. 2016), varying the constant effective population size (*N*_*e*_= 25,000 or 250,000) and the sample size (SS=10 or 20) and setting the mutation rate to *μ* = 10^-8. These sets of simulation parameters were combined with three values of the Block Penalty (BP) parameter of the LDhelmet program (see below), resulting in twelve conditions tested. For each combination of parameters, ten population samples were simulated, to generate independent replicates of the coalescent history (Figure 1A, Supplementary Figure S3).

Populations undergoing bottleneck and admixture events were also simulated with MSPRIME, using one of the five underlying landscapes. The sample size parameter was set to 20, and the *N*_*e*_ of the ancestral and present-day population was set to 250,000. The simulated bottleneck scenarios varied according to the timing of the bottleneck event (*t*_b_ = 500, 5,000, 50,000 generations ago) and the *N*_*e*_ of the population during the bottleneck (N_b_ = 2,500, 25,000). The duration of the bottleneck was fixed to 1,000 generations. The admixture scenarios varied in terms of the timing of the admixture event between the derived populations 1 and 2 (*t*_a_ = 500, 5,000, 50,000 generations ago). The time of split of the ancestral population (0) into two derived population 1 and 2 (*t*_split_) was set to 10*N_e_ generations. For each bottleneck and admixture scenario, ten replicated populations were simulated (Figure 1A,B).

A VCF file was generated with MSPRIME for each simulated population (Figure 1A, Supplementary Figure S3), which contains the genotypes of variants that segregate in the population sample consisting of 2*n* sequences (with *n* being the number of diploid samples) following the given underlying recombination landscape.

The impact of phasing errors on the inference of recombination rates was also assessed. Two replicate VCFs, simulated using the same underlying landscape, were manually dephased and then phased using Shapeit 4.2.2 (Delaneau et al., 2019). The phasing error rate was computed using the --switch-error option of VCFtools 0.1.17 (Danecek et al. 2011; using the original phased VCF as a reference). Phasing error rates of 2, 4, 6, 8 and 10% between heterozygous positions of the two original phased VCFs were then randomly generated, to produce five independent replicate VCF for each phasing error rate value.

### Inferred landscapes

Recombination rates were estimated for each of the simulated samples with LDhelmet (v1.19, Chan et al. 2012, Figure 1). Briefly, LDhemet uses phased sequence data to infer the ρ parameter locally, using likelihood computation between pairs of SNPs and then averaging over 50 consecutive variants to obtain a composite likelihood. The ρ parameter is inferred with a reversible-jump Markov Chain Monte Carlo algorithm using a step function applied to every window of 50 consecutive SNPs and determined by three parameters: the number of change-points, the locations of changes, and the recombination rate value of each constant fragment between two changes. We used VCFtools 0.1.17 (Danecek et al. 2011) and the vcf2fasta function of vcflib (https://github.com/vcflib/vcflib) to convert the SNP data obtained from MSPRIME simulations into the input format to LDhelmet, consisting of FASTA sequences of each individual haplotype. Ancestral states defined as the reference allele of each variant were also used as inputs. Each simulated replicate was analysed with LDhelmet using the following parameters. The haplotype configuration files were created with the find_conf function using the recommended window size of 50 SNPs. The likelihood look-up tables were created with the table_gen function using the recommended grid for the population recombination rate (*ρ*/pb) (*i*.*e. ρ* from 0 to 10 by increments of 0.1, then from 10 to 100 by increments of 1), and with the Watterson *θ* = 4*N*_*e*_*μ* parameter corresponding to the condition analysed. The Padé files were created using 11 Padé coefficients as recommended. The Monte Carlo Markov chain was run for 1 million iterations with a burn-in period of 100,000 and a window size of 50 SNPs. An important parameter to LDhelmet is the block penalty (BP), which determines the number of change-points, and thus the variance of the inferred recombination rates at a fine scale (*i*.*e*. smaller block penalty values correspond to a lower penalty for background rate changes, and thus generate more heterogeneous recombination landscapes). For each simulated combination of *N*_*e*_ and SS for populations with constant size, the block penalty was set to either 5, 10 or 50, and for the simulated populations undergoing bottleneck or admixture event, as for the “phase-error” datasets, the block penalty used was set to 5. Finally, the population recombination rates between each SNP pair were extracted with the post_to_text function, and were reported in ρ=4*N*_*e*_*r* per pb unit.

The reliability of the inferences was evaluated in various ways. For each combination of *N*_*e*_, SS, BP parameters and demographic scenarios simulated, the inferred, simulated and underlying landscapes were compared, in order to assess the ability of LDhelmet to reliably infer the true biological landscape (Figure 1, Supplementary Figure S3A, and see below Hotspot detection and Statistical Analysis). In order to evaluate the convergence of LDhelmet inferences across replicate runs, LDhelmet was run 10 times, for the 12 combinations of the parameters *N*_*e*_, SS and BP, on two independently simulated VCF files from constant-size populations sharing the same underlying landscape (Figure 1A, Supplementary Figure S3B). Finally, the inferred recombination landscapes of pairs of populations sharing the same underlying landscape were compared in order to assess the reproducibility of the LDhelmet inference, *i*.*e*., the expected variance between inferred maps in the absence of underlying biological variation (Figure 1, Supplementary Figure S3B).

### Variation in the μ/r ratio

To explore the influence of variation in mutation and recombination rates on the inference of recombination maps, two additional underlying landscapes were generated using the same procedure, this time targeting a ten times higher (i.e. 20 cM/Mb) or ten times lower (0.2 cM/Mb) mean recombination rate. Then, using one of the 5 underlying landscapes (*r* ∼ 10^-8 M/pb) and the 2 newly generated landscapes with mean *r* = 10^-7 and 10^-9 M/pb, respectively, sets of simulations were run with a *μ*/*r* ratio of 0.1, 1 and 10. This was achieved by fixing *μ* to either 10^-9, 10^-8 or 10^-7, while keeping a fixed *N*_*e*_ = 100000 and SS = 20 (Supplementary Table S1). For each of the tested combinations of *μ* and *r*, 10 populations were simulated. These simulated landscapes were inferred with LDhelmet, using a block penalty of 5.

### Hotspot detection

Underlying and simulated landscapes were first converted into population recombination rate landscapes by scaling them by 4*N*_*e*_. Underlying, simulated and inferred landscapes were then smoothed at a 500 bp and 2500 bp resolution using the Python package *scipy*.*stats*. The former corresponds to the underlying landscape resolution, and the latter to a trade off between the density of segregating sites and the resolution often used in the literature. For the different combinations of *N*_*e*_, SS and BP of the constant-size populations simulated with the 5 underlying landscapes, a mean simulated landscape and a mean inferred landscape were generated by averaging recombination rates across replicates.

Recombination hotspots of the underlying, simulated and inferred landscapes were called by comparing local vs surrounding recombination rates at each genomic window. A hotspot was defined as a window of 2.5 kb with an average recombination rate either 2.5, 5 or 10 times higher than the 50 kb flanking regions (excluding the focal window). Hotspot locations were then compared among landscapes using the same threshold values (Figure 1C)

### Statistical analyses

Statistical analyses were run with R 4.0.3. The length of underlying, simulated and inferred maps (*L*) was calculated at the 2.5 kb resolution using the formula:

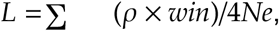

with *ρ* the population-scaled recombination rate, *win* the window size resolution used to smooth the maps in bp, and *N*_e_ the effective size of the simulated population. Several measures of the sensitivity, specificity, reliability, and repeatability of LDhelmet were computed, using the mean simulated and inferred landscapes of the constant-size populations, and replicates of simulated and inferred landscapes of the populations that underwent bottleneck of admixture events. Spearman rank correlation coefficients were calculated between the underlying and the corresponding simulated landscapes, between the simulated and inferred landscapes, and pairwise Spearman coefficients among the 10 replicates inferred from the two simulated populations sharing the same underlying landscape. True/false positive rates (TPR = TP/(TP+FN); FPR = FP/(FP+TN)), true/false negative rates (TNR = TN/(TN+FP); FNR = FN/(FN/(TP+FN)), and true/false discovery rates (TDR = TP/(TP+FP); FDR = FP/(TP+FP)) were calculated by comparing the simulated and inferred landscapes. The mean pairwise linear correlation (R^2^) and the proportion of shared hotspots was calculated between the 5 underlying landscapes, and for each condition simulated with a constant population size scenario, and for the three threshold values tested (*i*.*e*. 2.5, 5 and 10) between the simulated and inferred landscapes from the 5 different underlying landscapes, as well as between the pairs of populations sharing the same underlying landscape.

The statistical analyses were performed using home-made R scripts available upon request.

## Data, script and code availability

The underlying landscapes, the main scripts used to generate the underlying landscapes, run the simulations under the various demographic scenarios, infer the simulated landscapes with LDhelmet, and call hotspots from the landscapes can be found at https://doi.org/10.5281/zenodo.7657101. The Singularity container recipe built to run the simulations is available at: https://doi.org/10.5281/zenodo.7657199. This recipe contains the installation command lines of the required programs, the scripts used for the simulations, and the five underlying landscapes used in our study. More scripts and data are available upon request.

## Supplementary information

Supplementary Materials, containing Supplementary Figures and Tables are available at https://doi.org/10.5281/zenodo.7544555.

## Acknowledgments

We are grateful to Julien Joseph, Nicolas Lartillot, Frédéric Baudat, Bernard de Massy and Laurent Duret for the helpful discussions and feedback. We also thank Khalid Belkhir, Jimmy Lopez and Mathieu Massaviol for their support to build the Singularity container used to run our simulations.

