## Supplementary Materials for "Performance and limitations of linkage-disequilibrium-based methods for inferring the genomic landscape of recombination and detecting hotspots: a simulation study"

### Supplementary Figures

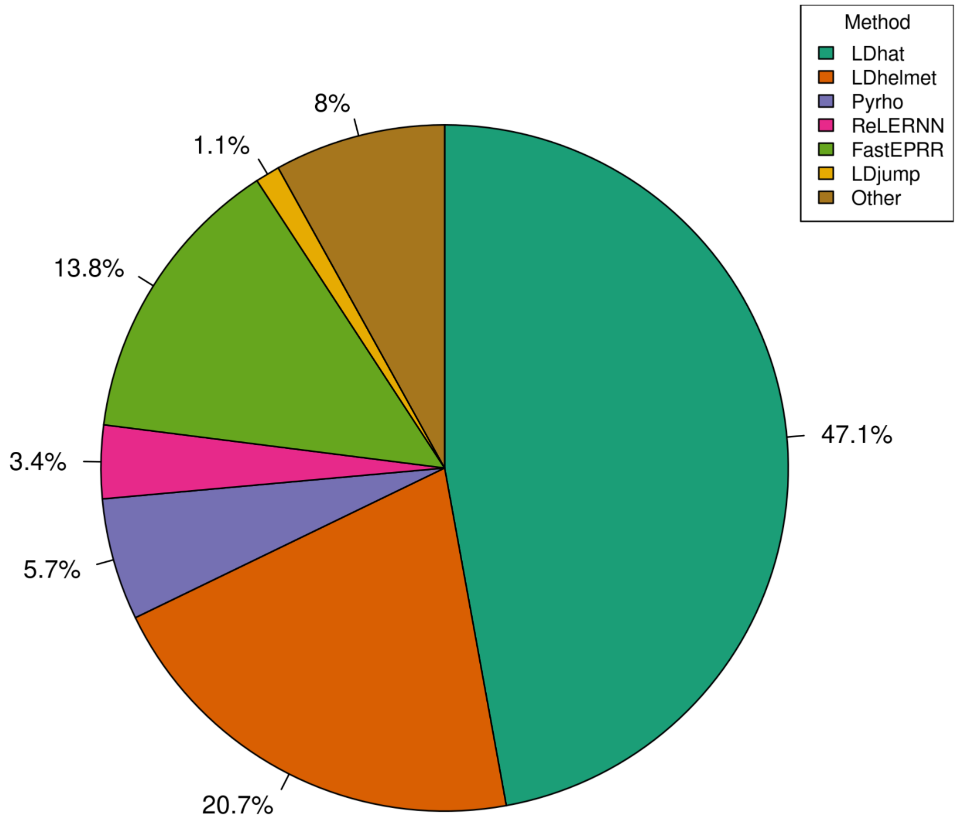


**Supplementary Figure S1.** Population-based methods used to infer recombination landscapes since 2007. Bibliography search for articles published in the last 15 years that built population-based recombination landscapes in plants, animals and bacteria. The search was first done using the keywords “population-based recombination landscape”, “linkage disequilibrium”, “hotspots”, “fine scale recombination landscape”; then retrieving all the papers that cite one of the methods identified to infer population-based recombination landscapes (i.e. LDhat, LDhelmet, Pyrho, ReLERNN, FastEPRR, LDjump, PHASE, iSMC, LDpop, RecMin, inferRho, heRho) and sorting the ones that used them to build recombination landscapes. The colors of the pie-chart shows the relative proportion of the use of the different methods in 87 articles identified to have built population recombination landscapes since 2007. The category “Other” contains the methods PHASE, iSMC, LDpop, RecMin and inferRho.

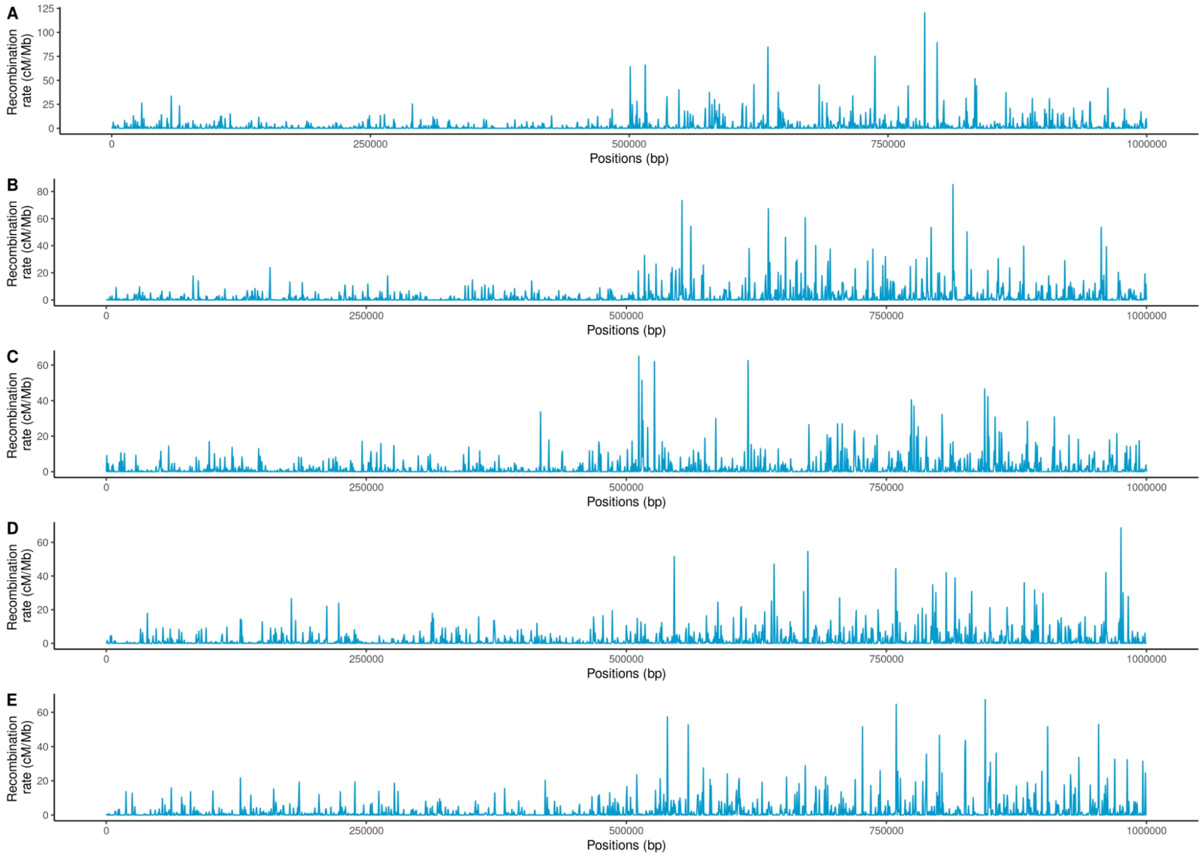


**Supplementary Figure S2.** The 5 underlying recombination landscapes (represented in units of cM/Mb (y-axis) along a chromosomal region of 1Mb (x axis)) generated using human ChIP-seq data from Pratto et al. (2014). The averaged recombination rate of the left half of the chromosome segment is 1 cM/Mb, and the right half is 3 cM/Mb.

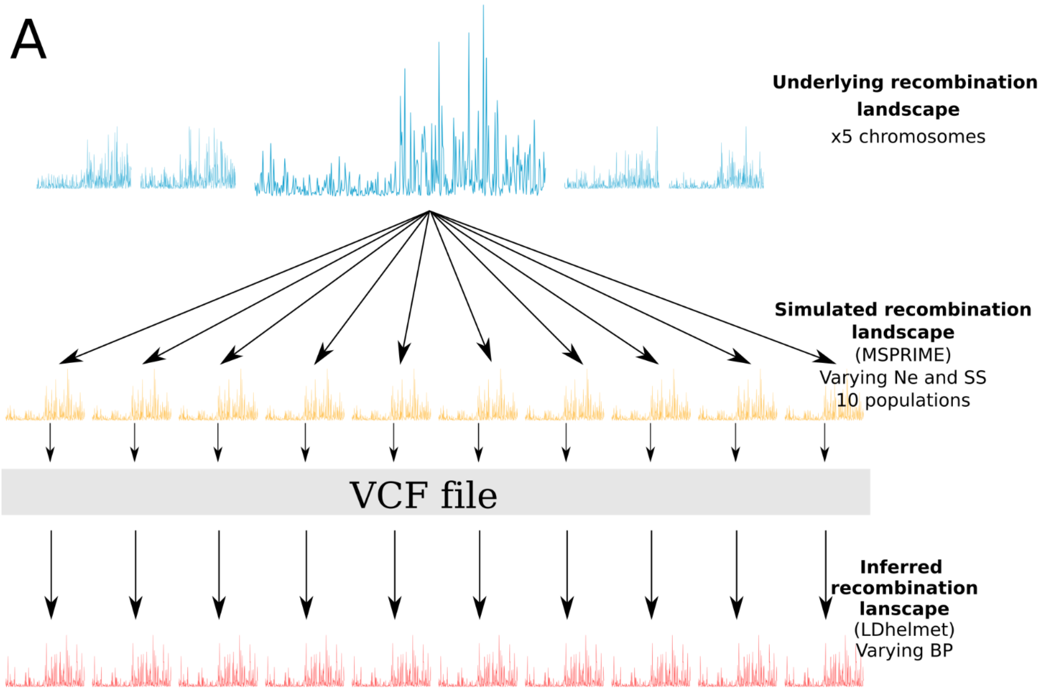


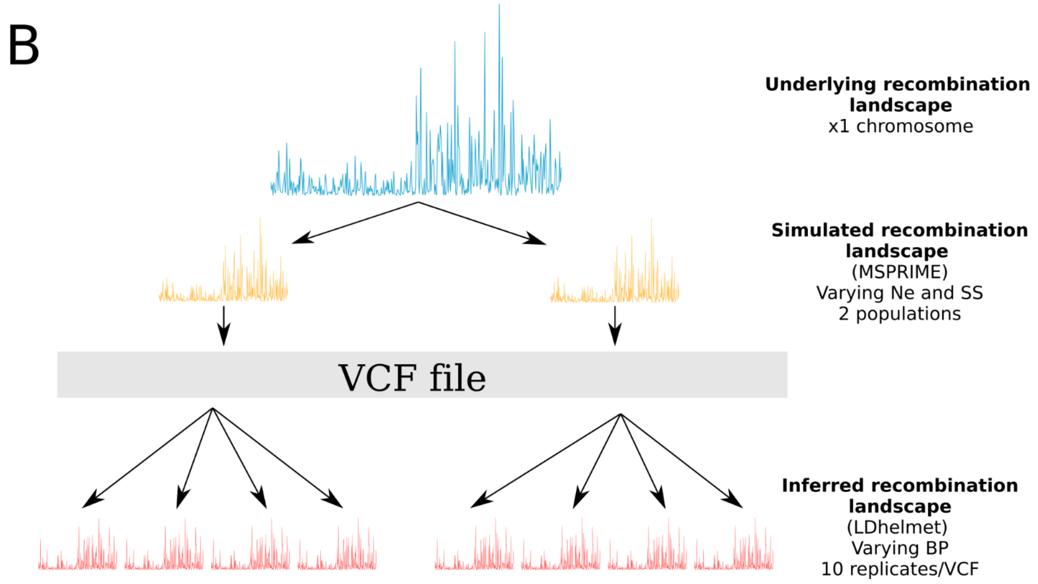


**Supplementary Figure S3.** Simulation and inference protocols with constant population size scenarios. **A)** In the first simulation framework, 5 different underlying recombination landscapes were generated based on human empirical data. These 5 landscapes can either be considered as parts of different chromosomes from the same species, or as orthologous parts of chromosomes from different species. For each of the 5 underlying landscapes, 10 recombination landscapes were simulated with MSPRIME 0.7.4 for 4 combinations of effective population size ($N_e$) and sample size (SS) parameters, generating a VCF file for each simulated population. The VCF files were then used to infer the local population recombination rates using LDhelmet 1.19 with 3 alternative block penalty values (BP, a key parameter to LDhelmet), representing a total of 12 tested conditions. **B)** In the second simulation framework, only one of the 5 underlying landscapes was used to generate 2 simulated populations for each of the 4 combinations of simulation parameters, before running LDhelmet 10 times in replicate for 3 different values of block penalty, representing a total of 12 tested conditions.

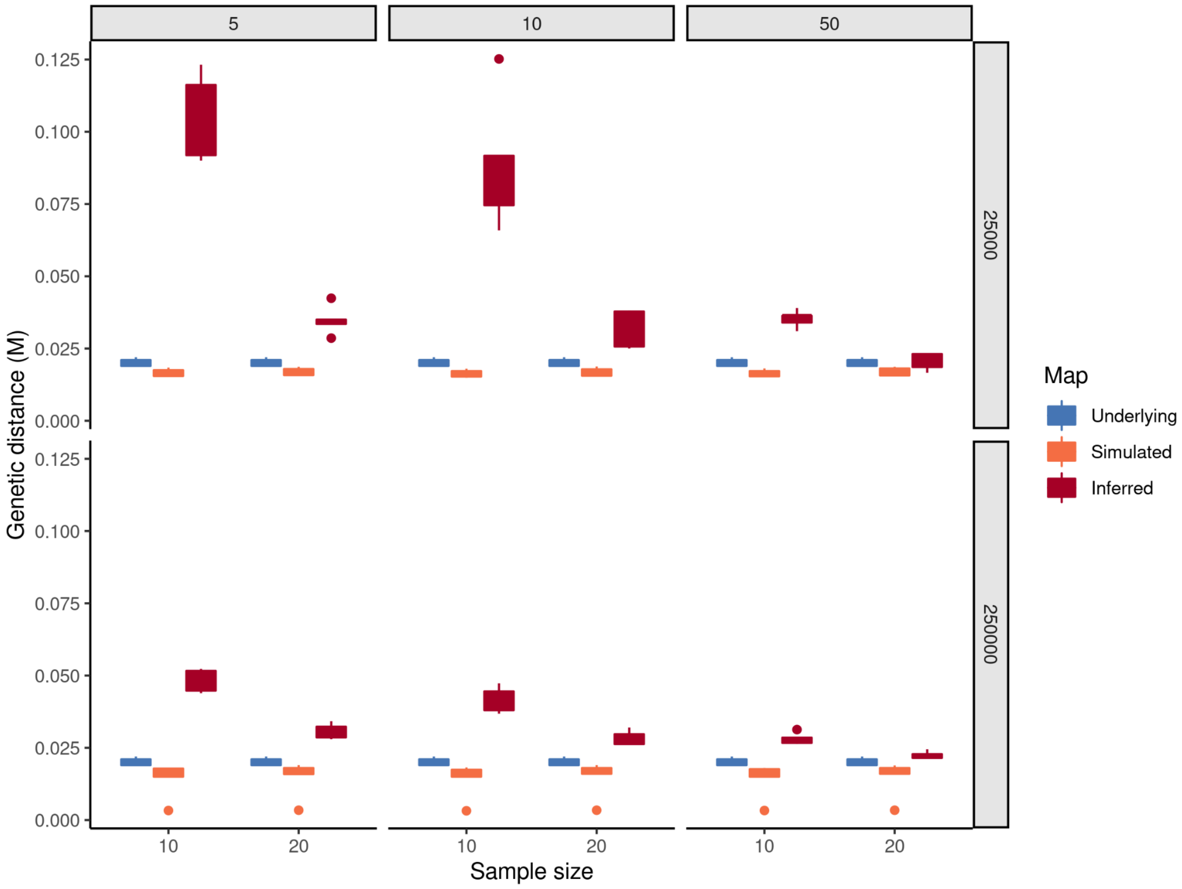


**Supplementary Figure S4**. Underlying (blue), simulated (orange) and inferred (red) recombination map length (in Morgan). The maps were smoothed at a 2.5 kb resolution.

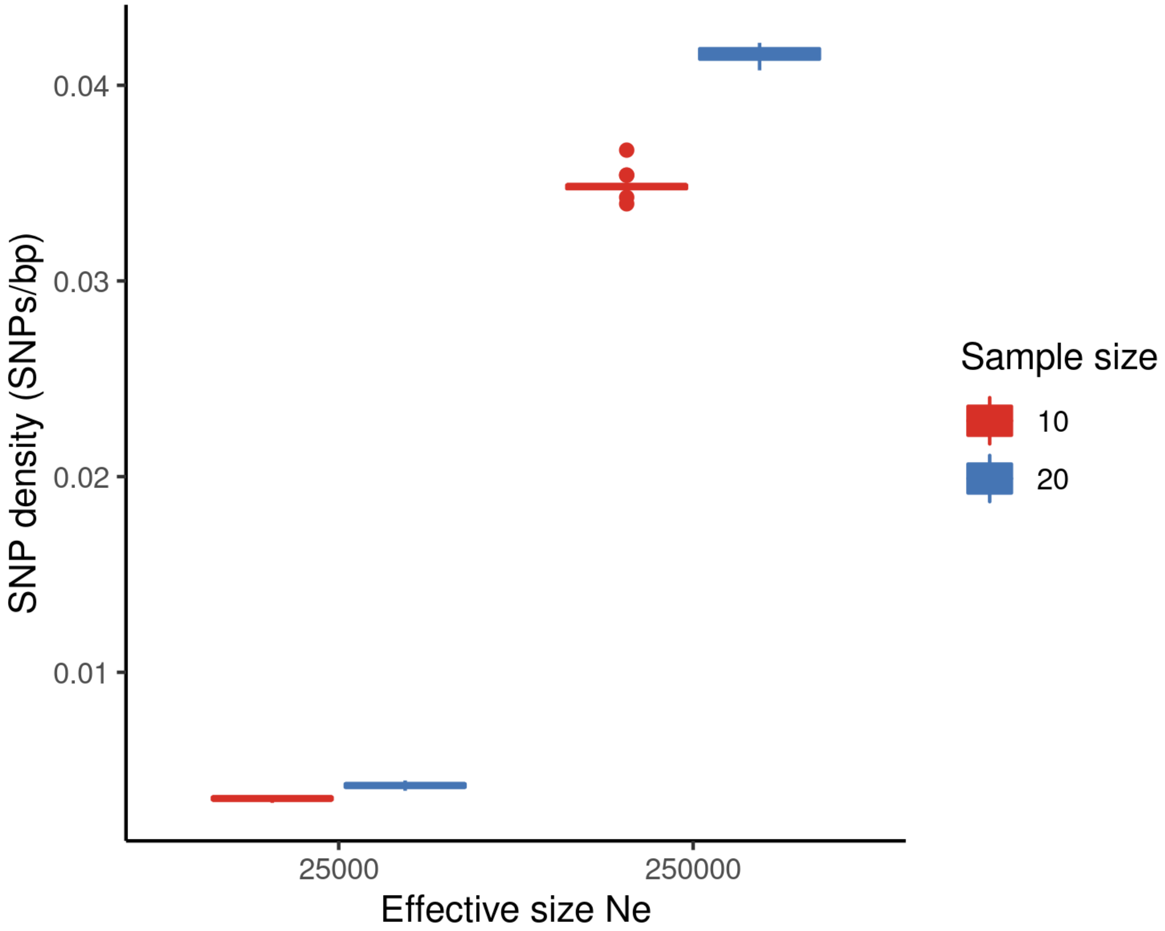


**Supplementary Figure S5**. SNP density according to $N_e$ and SS. The sample size (SS) parameter is shown in color (i.e. 10, blue or 20, red), and the effective population size ($N_e$) is shown on the x axis.

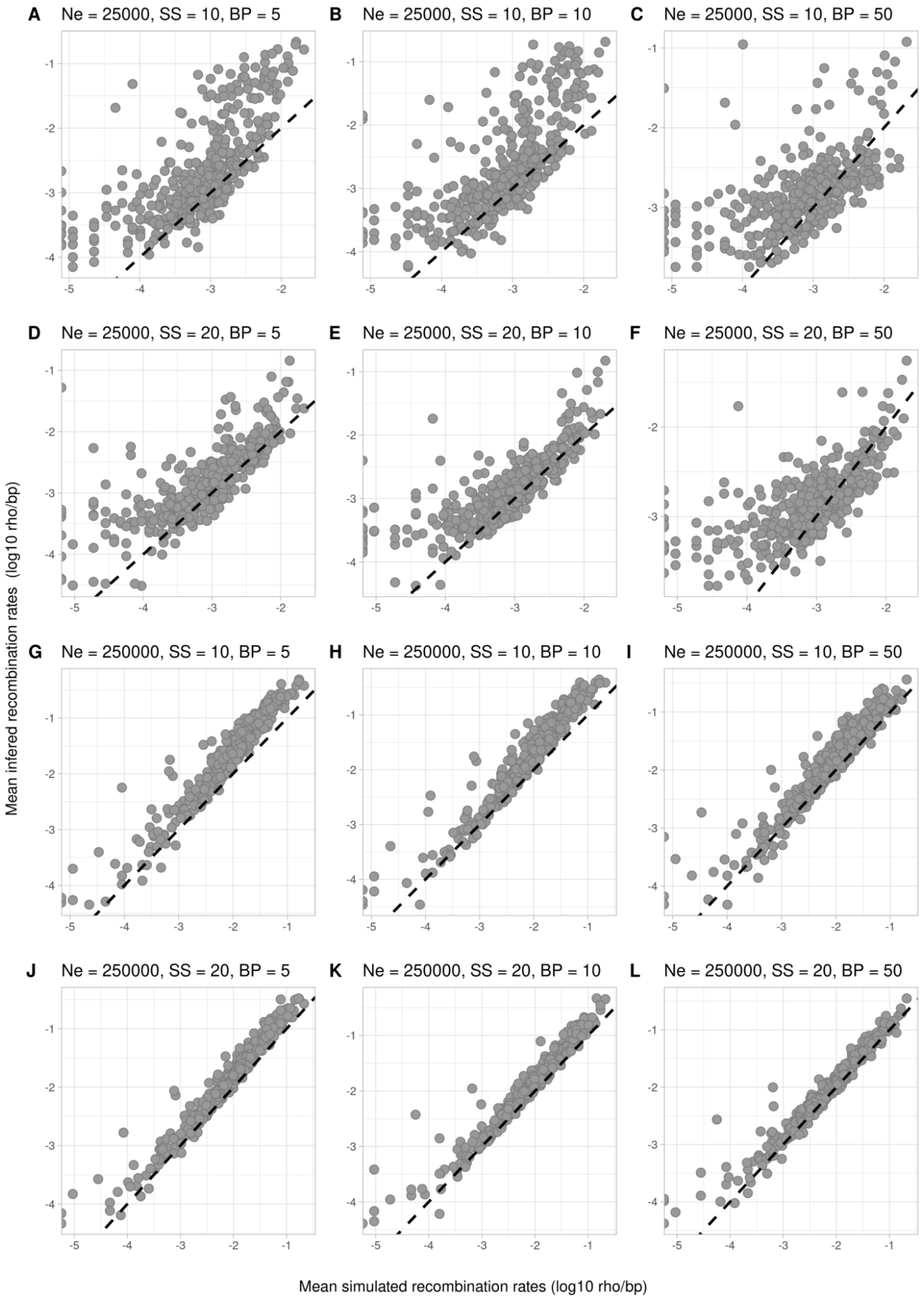


**Supplementary Figure S6.** Quality assessment of local recombination rates estimated by LDhelmet. Simulated and inferred recombination rates were averaged within 2.5kb windows across the replicates, for all the combination of parameters tested (i.e. $N_e$, SS, BP). The x axis shows the recombination rates of the mean simulated landscapes and the y axis the recombination rates of the mean inferred landscapes, both on a logarithmic scale. Each point corresponds to a local 2.5kb-window averaged across 10 replicated populations obtained under identical simulation parameters. The black dashed lines indicate the expectation of y=x. **A)** $N_e$ = 25,000, SS = 10, BP = 5. **B)** $N_e$ = 25,000, SS = 10, BP = 10. **C)** $N_e$ = 25,000, SS = 10, BP = 50. **D)** $N_e$ = 25,000, SS = 20, BP = 5. **E)** $N_e$ = 25,000, SS = 20, BP = 10. **F)** $N_e$ = 25,000, SS = 20, BP = 50. **G)** $N_e$ = 250,000, SS = 10, BP = 5. **H)** $N_e$ = 250,000, SS = 10, BP = 10. **I)** $N_e$ = 250,000, SS = 10, BP = 50. **J)** $N_e$ = 250,000, SS = 20, BP = 5. **K)** $N_e$ = 250,000, SS = 20, BP = 10. **L)** $N_e$ = 250,000, SS = 20, BP = 50.

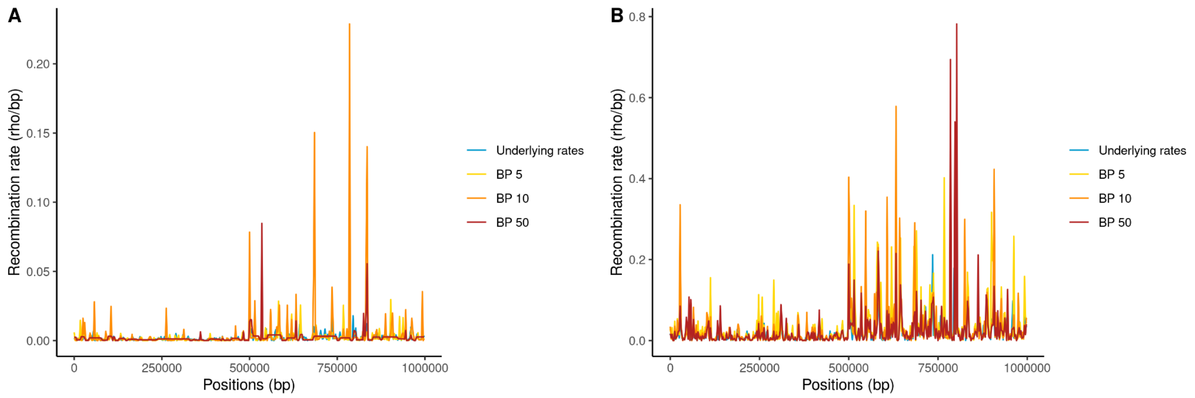


**Supplementary Figure S7.** Block penalty influences the recombination rate inferred by LDhelmet. The recombination landscape (in rho/bp) of one underlying landscape is shown in blue. The corresponding recombination landscape inferred with a BP = 5, 10 and 50 are shown in yellow, orange and red respectively, for SS = 20 and $N_e$ = 25000 (**A**) and 250000 (**B**).

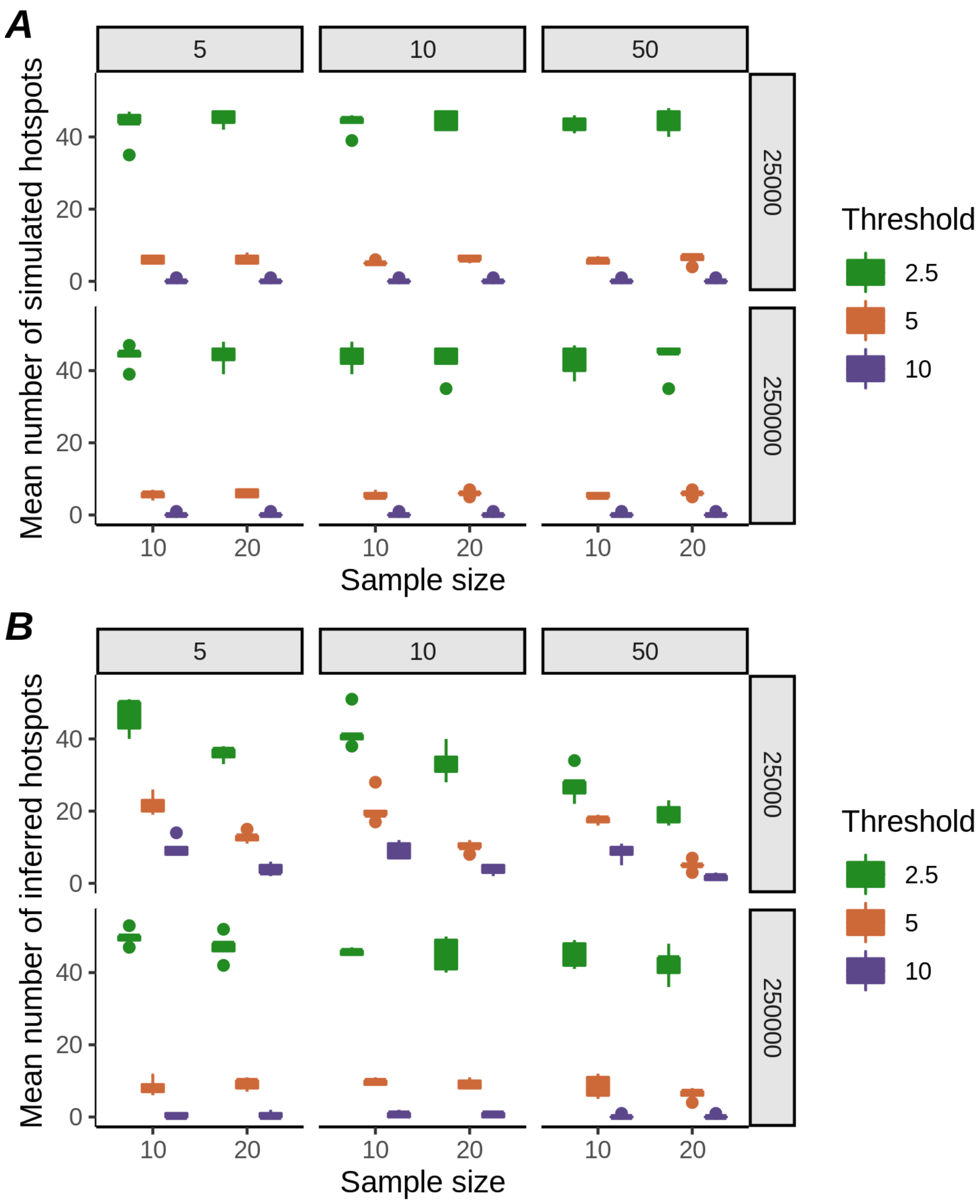


**Supplementary Figure S8**. Hotspot calling. Number of simulated (**A**) and inferred (**B**) hotspots called using three different threshold shown by colors (i.e. 2.5, 5 and 10) according to the parameters simulated (i.e. $N_e$, SS, BP). The sample size parameter is shown on the x axis (i.e. 10 or 20), the upper panels correspond to conditions where $N_e$ = 25 000, the lower panels correspond to conditions where $N_e$ = 250 000; and the BP parameters correspond to the vertical panels (i.e. 5, 10 and 50).

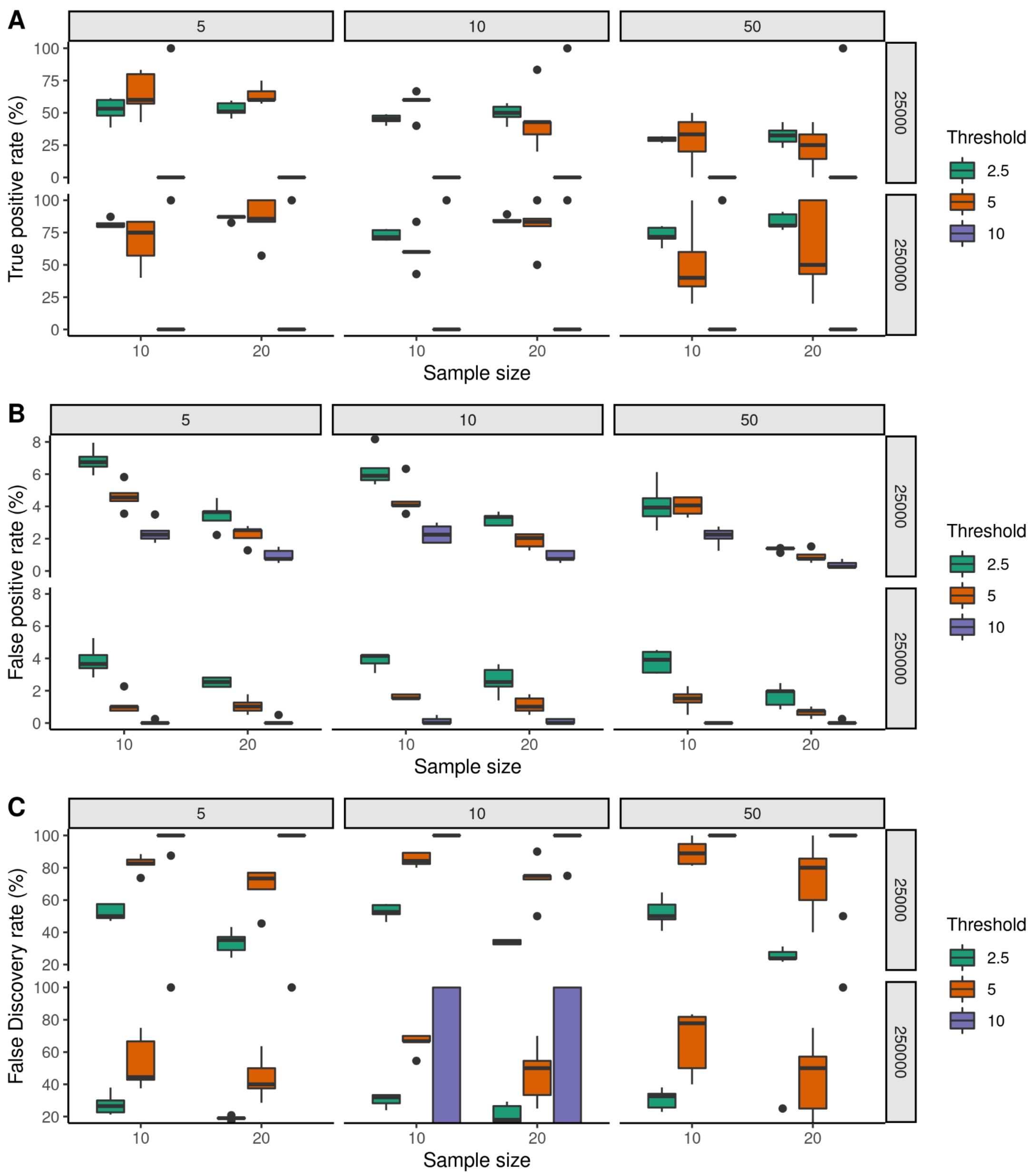


**Supplementary Figure S9.** Hotspot detection. True positive (TP, **A**), false positive (FP, **B**) and false discovery (FD, **C**) rates of inferred, as compared to simulated, hotspots, called from the averaged landscape using different threshold values (i.e. 2.5, 5, 10) shown in color, according to the parameters simulated. The sample size parameter is shown on the x axis (i.e. 10 or 20), the upper panels correspond to conditions where $N_e$ = 25,000, the lower panels correspond to conditions where $N_e$ = 250,000, and the BP parameter values correspond to the vertical panels (i.e. 5, 10 and 50).

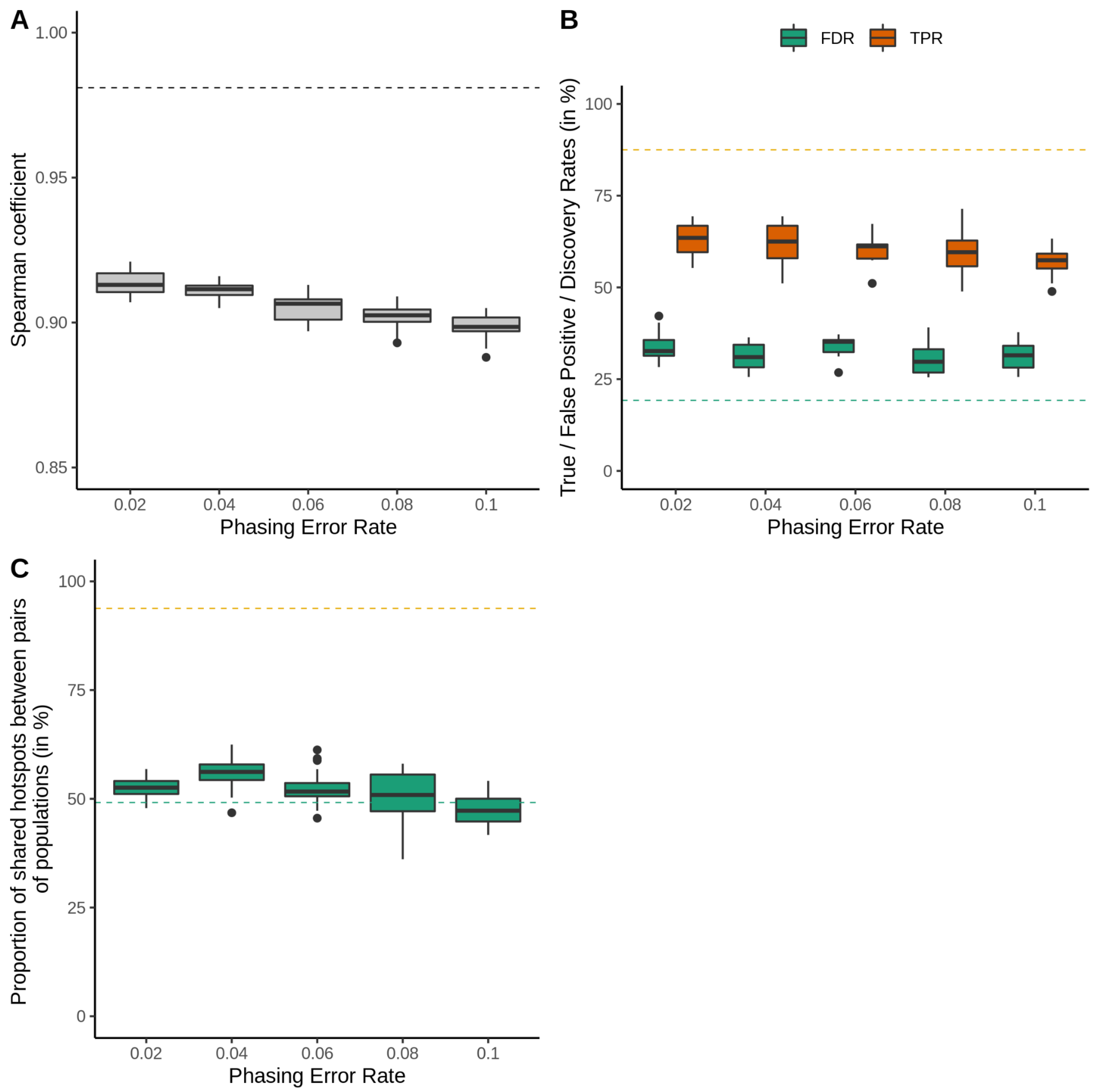


**Supplementary Figure S10.** Influence of phasing errors on recombination rate inference and hotspot detection. **A)** Spearman correlation coefficients between simulated and inferred landscapes; **B)** True positive (TP, orange) and false discovery (FD, green) rates of inferred, as compared to simulated, hotspots, called using a threshold of 2.5; and **C)** Proportion of shared hotspots between pairs of inferred (green) recombination landscapes, as compared to the proportion of shared hotspots between pairs of simulated landscapes in data without phasing errors (orange dashed line); according to the phasing error rate. Dashed lines in A, B, C) represent the corresponding values for simulations with truly phased data.

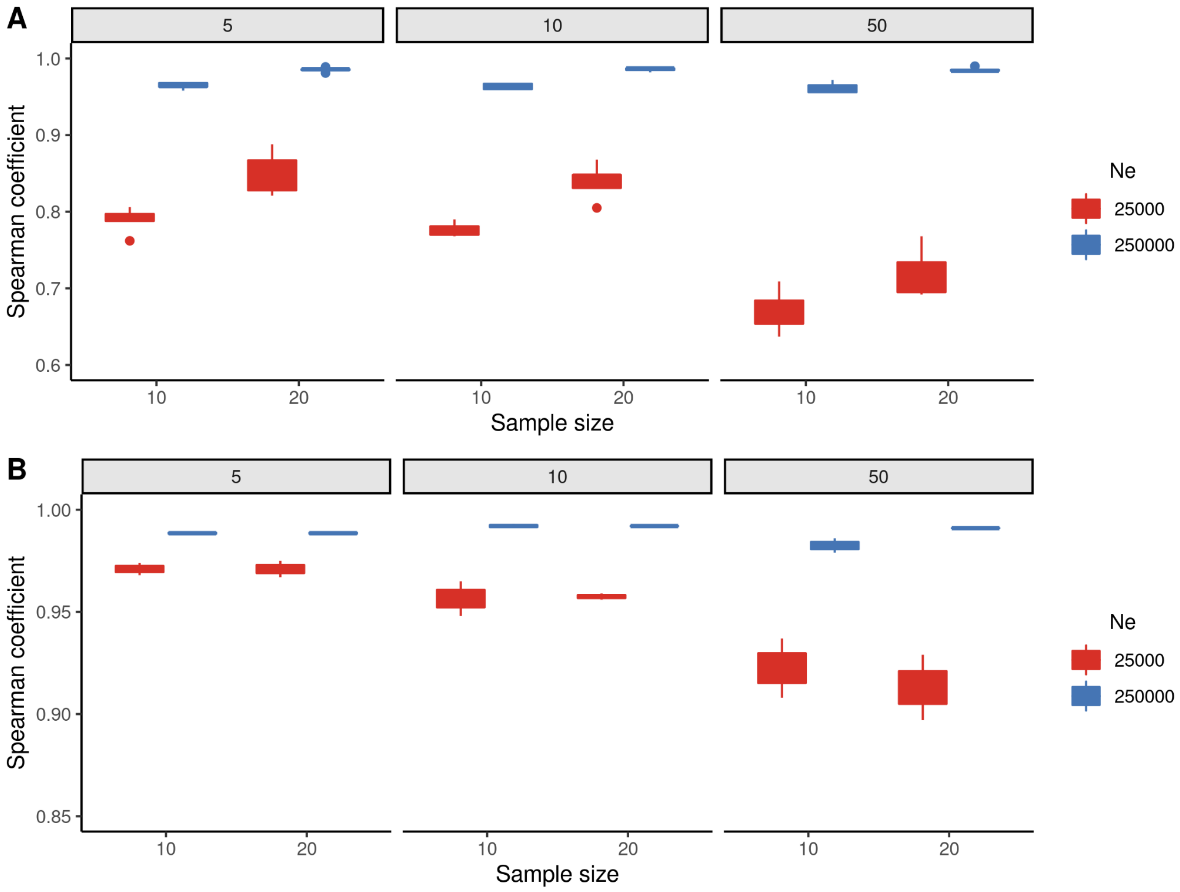


**Supplementary Figure S11.** Correlations between simulated and inferred landscapes (A), and between replicates of inferred landscapes (B) according to the differents parameters tested (i.e. $N_e$, SS, BP). The sample size is shown on the x axis (i.e. SS=10 or 20), the $N_e$ parameter is indicated by color (i.e. 25,000 in red and 250,000 in blue) and the LDhelmet BP values correspond to the different panels (i.e. BP=5, 10 or 50). **A)** Spearman correlation coefficients between the mean simulated and the mean inferred landscape calculated across the 10 replicated populations originating from each of the five different underlying landscapes (i.e. using simulation framework of Supplementary Figure S3A). **B)** Mean Spearman correlation coefficients calculated between pairwise comparisons among the ten replicates of inferred landscapes, from simulated populations sharing the same underlying landscape (i.e. using simulation framework of Supplementary Figure S3B).

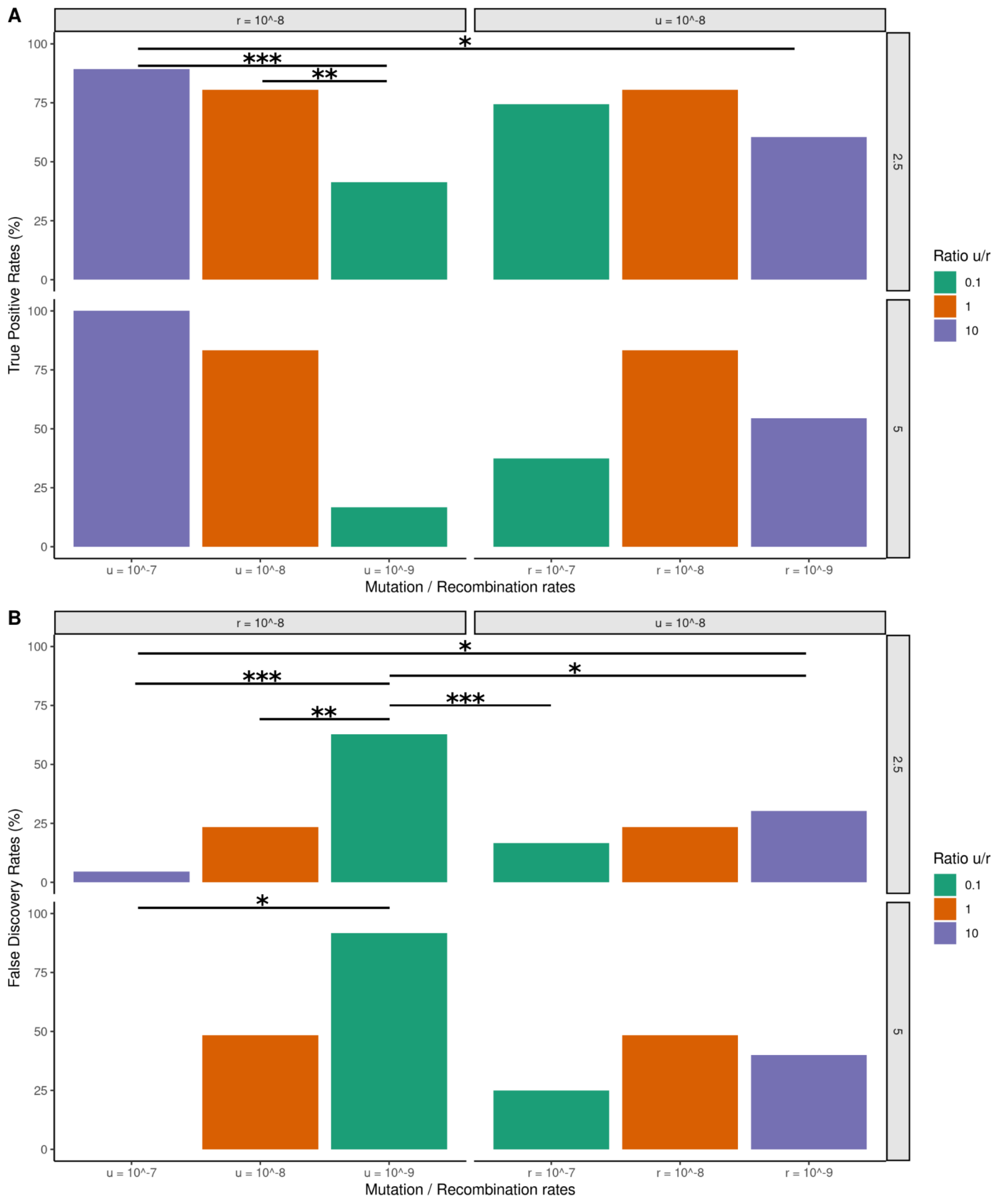


**Supplementary Figure S12.** Influence of the u/r ratio on hotspot detection. True positive (TP, **A**) and false discovery (FD, **B**) rates of hotspots, called using a threshold of 2.5 (upper panels) and 5 (lower panels) according to u and r. The x axis indicates u values when r=10^-8 (left panels), and r values for u=10^-8 (right panels). Colors correspond to the u/r ratio. The asterisks show to the prop.test p-value (* prop.test p-value<0.05, **<0.01, ***<0.001).

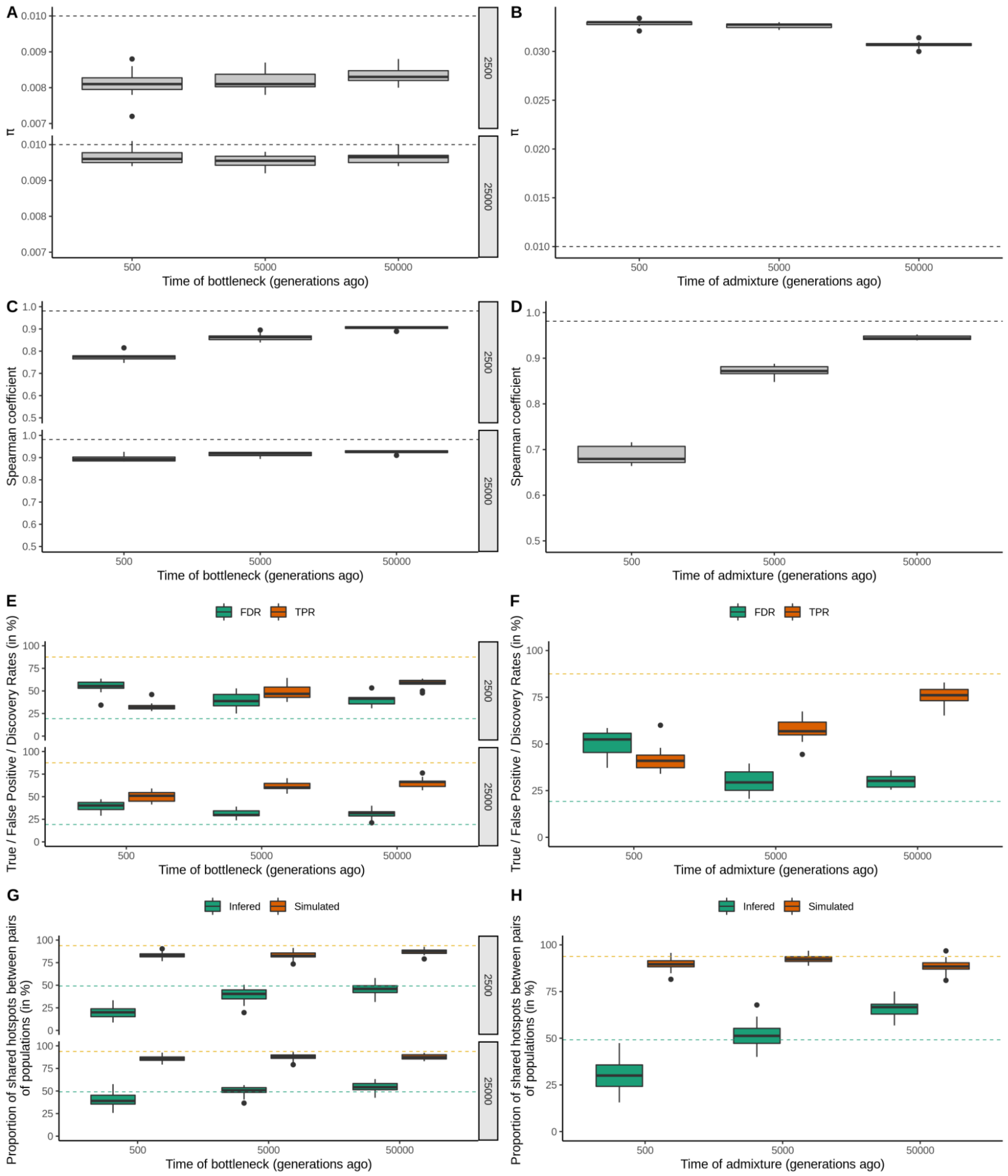


**Supplementary Figure S13.** Influence bottleneck and admixture events on recombination rate inference and hotspot detection. **A)** Nucleotide diversity, **B)** Spearman correlation coefficients between simulated and inferred landscape **C)** True positive (TP) rates and **D)** false discovery (FD) rates of inferred, as compared to simulated hotspots, called using a detection threshold of 2.5; according to the age and the strength ($N_b$, Figure 1) of the bottleneck for populations that underwent a bottleneck (left panels), and according to the age of the admixture event for the populations that underwent admixture (right panels). Results are shown for simulations with the parameters $N_e$=250000, SS=20 and BP=5.

### Supplementary Tables

| | | | | | Threshold = 2.5 | | Threshold = 5 | | Threshold = 10 | |
|---|---|---|---|---|---|---|---|---|---|---|
| Ratio | r | u | Correlation simulated-inferred | Mean pairwise correlation | TPR | FDR | TPR | FDR | TPR | FDR |
| **10** | 10-8 | 10-7 | 0.989 | 0.882 | 89.36 | 4.55 | 100.00 | 0.00 | 0 | 0 |
| **1** | 10-8 | 10-8 | 0.951 | 0.788 | 82.22 | 19.57 | 66.67 | 42.86 | 0 | 100 |
| **0.1** | 10-8 | 10-9 | 0.697 | 0.511 | 41.30 | 62.75 | 16.67 | 91.67 | 0 | 100 |
| **0.1** | 10-7 | 10-8 | 0.966 | 0.771 | 74.47 | 16.67 | 37.50 | 25.00 | 0 | 0 |
| **1** | 10-8 | 10-8 | 0.958 | 0.792 | 78.72 | 27.45 | 100.00 | 53.85 | 0 | 100 |
| **10** | 10-9 | 10-8 | 0.932 | 0.596 | 60.63 | 30.30 | 54.55 | 40.00 | 0 | 100 |

**Supplementary Table S1.** Influence of the *μ*/*r* ratio on recombination rate inference and hotspot detection. Spearman's correlation between mean simulated and mean inferred landscapes and averaged pairwise Spearman's correlation between the 10 inferred replicates are shown on the left half of the table. True positive (TP) and false discovery (FD) rates of hotspots called using three detection thresholds (i.e. 2.5, 5 and 10) are indicated in the right half of the table. Rows correspond to the different combinations of *μ* and *r* tested.

| | | Mean proportion of shared simulated hotspots | | | | Mean proportion of shared inferred hotspots | | |
|---|---|---|---|---|---|---|---|---|
| | $R^2$ log(simuli~simulj) | Threshold = 5 | Threshold = 2.5 | Threshold = 10 | $R^2$ log(inferi~inferj) | Threshold = 5 | Threshold = 2.5 | Threshold = 10 |
| **Different underlying landscapes** | | | | | | | | |
| Ne = 25000 SS = 10 BP = 5 | 0.012 | 1.715 | 8.065 | 0.00 | 0.033 | 5.385 | 13.99 | 3.215 |
| Ne = 25000 SS = 10 BP = 10 | 0.015 | 1.835 | 7.275 | 0.00 | 0.043 | 4.945 | 11.28 | 0.00 |
| Ne = 25000 SS = 10 BP = 50 | 0.014 | 1.55 | 7.595 | 0.00 | 0.039 | 5.075 | 5.895 | 2.51 |
| Ne = 25000 SS = 20 BP = 5 | 0.014 | 2.00 | 9.045 | 0.00 | 0.032 | 3.95 | 9.735 | 1.835 |
| Ne = 25000 SS = 20 BP = 10 | 0.013 | 1.55 | 8.505 | 0.00 | 0.042 | 1.125 | 7.46 | 0.00 |
| Ne = 25000 SS = 20 BP = 50 | 0.015 | 1.43 | 8.55 | 0.00 | 0.084 | 0.00 | 4.28 | 0.00 |
| Ne = 250000 SS = 10 BP = 5 | 0.017 | 2.085 | 8.42 | 0.00 | 0.013 | 0.00 | 11.245 | 0.00 |
| Ne = 250000 SS = 10 BP = 10 | 0.016 | 2.00 | 8.18 | 0.00 | 0.013 | 3.055 | 11.125 | 0.00 |
| Ne = 250000 SS = 10 BP = 50 | 0.015 | 2.00 | 7.11 | 0.00 | 0.012 | 0.97 | 9.86 | 0.00 |
| Ne = 250000 SS = 20 BP = 5 | 0.017 | 1.715 | 7.675 | 0.00 | 0.014 | 1.17 | 9.55 | 0.00 |
| Ne = 250000 SS = 20 BP = 10 | 0.013 | 1.55 | 7.12 | 0.00 | 0.014 | 1.18 | 8.695 | 0.00 |
| Ne = 250000 SS = 20 BP = 50 | 0.016 | 1.67 | 8.075 | 0.00 | 0.015 | 1.34 | 7.32 | 0.00 |
| **Same underlying landscape** | | | | | | | | |
| Ne = 25000 SS = 10 BP = 5 | 0.7463 | 85.71 | 79.62 | 0 | 0.2139 | 30.65 | 28.55 | 20.95 |
| Ne = 25000 SS = 10 BP = 10 | 0.7386 | 72.92 | 77.50 | 0 | 0.1429 | 10.9 | 12.9 | 0 |
| Ne = 25000 SS = 10 BP = 50 | 0.7101 | 77.78 | 71.15 | 0 | 0.1173 | 10.7 | 21.7 | 0 |
| Ne = 25000 SS = 20 BP = 5 | 0.7497 | 80.36 | 78.99 | 0 | 0.311 | 14.85 | 26.9 | 0 |
| Ne = 25000 SS = 20 BP = 10 | 0.7908 | 78.75 | 72.61 | 0 | 0.3035 | 27.7 | 23.55 | 22.9 |
| Ne = 25000 SS = 20 BP = 50 | 0.7622 | 65.00 | 76.09 | 0 | 0.2206 | 0 | 21.15 | 0 |
| Ne = 250000 SS = 10 BP = 5 | 0.9466 | 80.36 | 90.42 | 0 | 0.6301 | 15.3 | 35.65 | 0 |
| Ne = 250000 SS = 10 BP = 10 | 0.9346 | 85.71 | 85.81 | 0 | 0.6671 | 31.75 | 38.95 | 0 |
| Ne = 250000 SS = 10 BP = 50 | 0.9391 | 81.25 | 90.33 | 0 | 0.6019 | 6.3 | 36 | 0 |
| Ne = 250000 SS = 20 BP = 5 | 0.9554 | 80.36 | 93.79 | 0 | 0.7514 | 32.55 | 49.15 | 41.65 |
| Ne = 250000 SS = 20 BP = 10 | 0.9562 | 92.86 | 92.57 | 0 | 0.7157 | 26.7 | 55.55 | 0 |
| Ne = 250000 SS = 20 BP = 50 | 0.9488 | 87.50 | 82.38 | 0 | 0.7421 | 56.9 | 54.25 | 0 |

**Supplementary Table S2.** Expected and observed hotspot sharing between populations with different or identical underlying landscapes. $R^2$ and mean proportion of shared hotspots between pairs of simulated (expected proportion, left half of the table) and pairs of inferred (observed proportion, right of the table) recombination landscapes of populations simulated with the same set of parameters (indicated by rows) and either sharing different underlying landscapes (top, following simulation framework from Supplementary Figure S3A), or identical underlying landscape (down, following simulation framework from Supplementary Figure S3B), for all combination of parameters tested (i.e. $N_e$, SS, BP).
